# A multi-omic integrative scheme characterizes tissues of action at loci associated with type 2 diabetes

**DOI:** 10.1101/2020.06.25.169706

**Authors:** Jason M. Torres, Moustafa Abdalla, Anthony Payne, Juan Fernandez-Tajes, Matthias Thurner, Vibe Nylander, Anna L. Gloyn, Anubha Mahajan, Mark I. McCarthy

**Affiliations:** The Wellcome Centre for Human Genetics, Nuffield Department of Medicine, University of Oxford, Oxford, OX3 7BN, UK; Oxford Centre for Diabetes, Endocrinology and Metabolism, Radcliffe Department of Medicine, University of Oxford, Oxford, OX3 7LE, UK; Division of Endocrinology, Department of Pediatrics, Stanford School of Medicine, Stanford, CA 94305, USA; Clinical Trial Service Unit and Epidemiological Studies Unit, Nuffield Department of Population Health, Big Data Institute, University of Oxford, Oxford, OX3 7LF, UK; OMNI Human Genetics, Genentech, 1 DNA Way, South San Francisco, CA 94080, USA

**Author notes:** These authors co-supervised this work (Anubha Mahajan, Mark I. McCarthy).

## Abstract

Resolving the molecular processes that mediate genetic risk remains a challenge as most disease-associated variants are non-coding and functional and bioinformatic characterization of these signals requires knowledge of the specific tissues and cell-types in which they operate. To address this challenge, we developed a framework for integrating tissue-specific gene expression and epigenomic maps (primarily from tissues involved in insulin secretion and action) to obtain *tissue-of-action* (TOA) scores for each association signal by systematically partitioning posterior probabilities from Bayesian fine-mapping. We applied this scheme to credible set variants for 380 association signals from a recent GWAS meta-analysis of type 2 diabetes (T2D) in Europeans. The resulting tissue profiles underscored a predominant role for pancreatic islets and, to a lesser extent, subcutaneous adipose and liver, that was largely attributable to enhancer elements and transcribed regions, particularly among signals with greater fine-mapping resolution. We incorporated resulting TOA scores into a rule-based classifier, and validated the tissue assignments through comparison with data from *cis*-eQTL enrichment, functional fine-mapping, RNA co-expression, and patterns of physiological association. In addition to implicating signals with a single tissue-of-action, we also found evidence for signals with shared effects in multiple tissues as well as distinct tissue profiles between independent signals within heterogeneous loci. Lastly, we demonstrated that TOA scores can be directly coupled with eQTL colocalization to further resolve effector transcripts at T2D signals. This framework guides mechanistic inference by directing functional validation studies to the most relevant tissues and can gain power as fine-mapping resolution and cell-specific annotations become richer. This method is generalizable to all complex traits with relevant annotation data and is made available as an R package.

## Introduction

The scale of genetic studies of type 2 diabetes (T2D) has dramatically expanded in recent years to encompass hundreds of thousands of individuals and tens of millions of variants, culminating in the discovery of over 400 independent genetic associations that influence disease susceptibility^1–3^. However, as with other complex traits, the majority of T2D-associated variants are non-coding and are presumed to mediate risk by affecting genetic regulatory mechanisms^4^. Characterization of the processes mediating genetic risk requires definition of the regulatory elements perturbed by these variants, along with the downstream consequences on gene expression and molecular pathways. Such regulatory insights have been typically gleaned through genome-wide approaches that integrate genetic data with information from expression quantitative trait loci (eQTL) analyses, chromatin accessibility and interaction mapping, and functional screening^5–10^.

A major challenge to these approaches is that the molecular processes that underpin disease risk are often tissue specific. Although the methods mentioned above can inform a genome-wide view of the tissues most prominently involved in disease (e.g. through patterns of genome-wide enrichment), they do not necessarily identify the most relevant tissue at any given association signal. For example, although several studies have shown strong enrichment of T2D-associated SNPs among regulatory elements in pancreatic islet tissue, there are clearly some signals that exert their impact on disease risk in peripheral tissues such as adipose, skeletal muscle, and liver^11–14^. Basing functional interpretation on the wrong tissue for a given variant (e.g. relying on islet data for a signal that operates in the liver) is likely to give rise to misleading inference and misdirected efforts at subsequent experimental characterization. Furthermore, as more detailed maps of regulatory elements and functional data in tissues and cell-types relevant to disease become available, the need to formulate principled strategies for integrating these features across datasets becomes more important, as the ever expanding scope of epigenomic and transcriptomic reference data can otherwise complicate variant interpretation.

To address the challenge of determining most likely *tissues-of-action* at loci associated with complex traits such as T2D, we developed a framework for jointly integrating genetic fine-mapping, gene expression, and epigenome maps across multiple disease-relevant tissues. As an illustration, we show how this scheme enabled a scalable approach for comparing the relative contributions of the key tissues involved in T2D pathogenesis (i.e. those controlling insulin secretion and action) by allowing us to delineate probabilistic tissue scores at individual genetic signals (deemed *tissue-of-action* or TOA scores). We explored the utility of this approach by applying it to a set of fine-mapped genetic associations from a recent large-scale meta-analysis of T2D and assessed the extent to which assigned tissues from a score-based classifier were corroborated by orthogonal datasets. We present results from these analyses along with new insights gleaned from specific loci that show, collectively, that this systematic approach to integrating disparate sources of information effectively resolves relevant tissues at GWAS loci.

## Methods

### Genetic data

Genome-wide association summary statistics from a meta-analysis of T2D GWAS corresponding to 32 studies of European ancestry (74,124 cases and 824,006 controls)^3^, conducted by DIAMANTE consortium, are available on the DIAbetes Genetics Replication And Meta-analysis (DIAGRAM) Consortium website (https://www.diagram-consortium.org). We used the summary statistics from the inverse-variance weighted fixed-effects meta-analysis of T2D-unadjusted for BMI that was corrected for residual inflation (accounting for structure between studies) with genomic control^3^. Of the 403 conditionally independent GWAS signals reported in Mahajan et al. 2018b, 380 signals were amenable to fine-mapping after excluding rare variants (e.g. minor allele frequency (MAF)<0.25%) and a signal mapping to the MHC locus^3^. The 99% genetic credible sets that corresponded to each signal and comprised SNPs that were each assigned a posterior probability of association (PPA) - summarizing the causal evidence for each SNP^15, 16^ - were also downloaded from the DIAGRAM website.

### Gene expression data

Gene expression data for 53 tissues - including liver, skeletal muscle, and subcutaneous adipose tissue - were downloaded from the Genotype-Tissue Expression Project (GTEx) Portal website (https://gtexportal.org). Data correspond to GTEx version 7 (dbGaP Accession phs000424.v7.p2) and represent RNA sequencing reads mapped to GENCODE (v19) genes^17^.

Gene expression data for pancreatic islets (n=114) was accessed from a previous study^5^ that involved sequencing stranded and unstranded RNA library preparations at the Oxford Genomics Centre. This set of islet samples was used to calculate expression specificity scores and perform coexpression analysis (see below and in the section titled “Gene co-expression”). An additional set of 60 islet samples available to us in-house were also used for eQTL mapping and enrichment analysis. All 174 islet samples were included in a subsequent analysis^18^ performed by the Integrated Network for Systemic analysis of Pancreatic Islet RNA Expression (InsPIRE) consortium. RNA-sequencing reads of all islet samples were also mapped to gene annotations in GENCODE (v19), in line with GTEx accessed data, using Spliced Transcripts Alignment to a Reference (STAR; v 020201) and quantified with featureCounts (v 1.50.0-p2).

Gene read counts for each tissue were transcript per million (TPM) normalised to correct for differences in gene length and library depth across samples. The tissue specificity of TPM-normalized gene expression was measured with expression specificity scores (ESS) obtained using the formula:

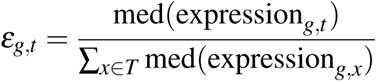

where *ε*_*g,t*_ is the ESS score for gene *g* in tissue *t*, and *T* is the set of evaluated tissues.

### Partitioning chromatin states

Chromatin state maps from a previous study^19^ based on a 13-state ChromHMM^20^ model trained from ChIP-seq input for histone modifications (H3K27ac, H3K27me3, H3K36me3, H3K4me1, and H3K4me3) were downloaded from the Parker lab website (https://theparkerlab.med.umich.edu). Chromatin state maps for liver, pancreatic islet, skeletal muscle, and subcutaneous adipose were used for the present study. Partitioned chromatin state maps used for generating tissue-of-action scores (see below in section titled “Deriving tissue-of-action (TOA) scores”), were obtained in the R statistical environment (v 3.6.0) using the Genomic Ranges (v 1.36.1) library. For each chromatin state annotation, the `disjoin` function (Genomic Ranges) was used to delineate non-overlapping segments across each of the four tissues. These segments were then compared with the annotation sets corresponding to each tissue to determine segments that were: (i) tissue-specific; (ii) shared across all tissues; or (iii) shared in a combination of two or more (but not all) tissues.

### Annotation enrichment analysis

To obtain fold enrichment values to use as annotation weights, genome-wide enrichment analysis was performed using the program fgwas^21^ (v 0.3.6), taking as input summary statistics from the DIAMANTE European BMI-unadjusted meta-analysis of T2D GWAS^3^. Enrichment of T2D-associated SNPs was assessed for coding sequence (CDS) and 13 chromatin state annotations mapped in human islet, liver, skeletal muscle, and subcutaneous adipose tissue from the Varshney et al. study^19^. To estimate log2-fold enrichment values, the `–cc` flag was used (specifying GWAS input from a case-control study) and default distance parameters were applied (i.e. genome partitioned ‘blocks’ of 5,000 SNPs). Weights were obtained by exponentiating the mean log2-fold enrichment values for each tissue-level annotation.

### Deriving *tissue-of-action* (TOA) scores

In order to obtain TOA scores for each of the 380 conditionally independent genetic association signals, we partitioned the corresponding PPA values of the 99% genetic credible set SNPs. For each SNP *j* in the 99% credible set, we obtain a vector *s* _*j,a*_ for each annotation *a* among the set of coding sequence and chromatin state annotations in set *A*. Each element in *s* _*j,a*_ corresponds to a tissue *t* in the set *T* comprising all evaluated tissues and is given by the equation:

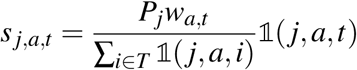

where *P*_*j*_ is the PPA of SNP *j, w*_*a,t*_ is the weight of annotation *a* in tissue *t*, and 1 is an indicator function defined as:

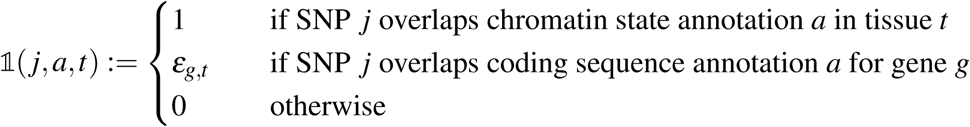

where *ε*_*g,t*_ is the ESS value for gene *g* in tissue *t*. Note that ESS values were used for coding SNPs as the relative expression levels of the corresponding gene can be used to inform tissue-level relevance for each coding SNP. If the SNP *j* does not map to annotation *a* in any tissue *t* ∈ *T*, the value of *s* _*j,a,t*_ is equated to 0. The vector *s* _*j*_ is thus given by:

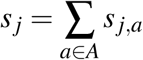

where the elements in *s* _*j*_ correspond to each tissue *t* ∈ *T* obtained from the linear combination of the partitioned PPA values across all annotations and weighted by the genome-wide fold enrichment of trait-associated variants for each tissue-level annotation. The vector *τ*_*c*_ that comprises TOA scores for each tissue *t* ∈ *T* and corresponds to 99% genetic credible set *c* is given by:

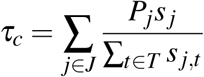

where *J* is the set of SNPs in the 99% genetic credible set *c*. Lastly, an unclassified score *U*_*c*_ is defined for each 99% genetic credible set *c*:

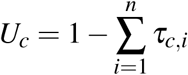

and indicates the cumulative PPA in *c* that is attributable to credible SNPs that do not map to any of the evaluated tissue-level annotations.

To evaluate the robustness of TOA score-based estimates of overall tissue contributions to T2D risk against the effect of GWAS association strength, we constructed weighted TOA scores:

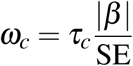

where *β* and SE are the effect size and standard error for the conditionally-independent SNP upon which the 99% credible set *c* was mapped.

### Profiling tissue specificity

The sum of squared distances (SSD) between TOA scores in *τ*_*c*_ for each *c* ∈ *C* (where *C* is the set of 99% genetic credible sets) was used as a measure of tissue specificity. To gauge the relationship between fine-mapping resolution and tissue-specificity, univariate linear models were used to estimate *β* coefficients corresponding to the regression of SSD on either the maximum 99% genetic credible set PPA, or the log10 number of SNPs in the 99% genetic credible sets. Signals were designated as “shared” if the difference between the top two TOA scores was ≤ 0.10. “Shared” signals were then tiered based on fine-mapping resolution: (i) Signals corresponded to 99% genetic credible sets comprised of a single credible SNP; (ii) Signals corresponded to 99% genetic credible sets where the maximum PPA ≥ 0.50 (i.e. where a single SNP explained most of the cumulative PPA); (iii) Signals corresponded to 99% genetic credible sets where the maximum PPA *<* 0.50. The relationship between SSD and fine-mapping resolution (i.e. maximum credible set PPA and number of credible SNPs) was visualized using the `scatterpie` library (v 0.1.4) in the R statistical environment (v 3.6.0).

### Rule-based classifier

A rule-based classifier for assigning each genetic signal (i.e. 99% genetic credible set) to a tissue was derived by assigning each genetic signal *c* to a tissue *t* if the corresponding TOA score in *τ*_*c*_ had the maximum value and exceeded a specified threshold. Sets of tissue-assigned signals were constructed for each stringency threshold within the set 0.0, 0.2, 0.5, 0.8. The classifier also allowed for a “shared” designation using the criteria described in the previous section (i.e. difference between the top two TOA scores was ≤ 0.10).

### eQTL mapping and tissue-specific eQTL enrichment

eQTLs for human liver, skeletal muscle, and subcutaneous adipose tissue were accessed from the GTEx Portal website (https://gtexportal.org) and corresponded to GTEx version 7 (dbGaP Accession phs000424.v7.p2). For human islet tissue, we used 174 samples (described above in section “Gene expression data”), and performed eQTL mapping using FastQTL (v 2.0) using a nominal pass with the –normal flag (to fit TPM-normalised read counts to a normal distribution). Gender and the first 15 PEER factors^22^ were used as covariates. For each tissue, q-values were calculated from nominal p-values and a false discovery rate threshold of ≤ 0.05 was applied to identify significant eQTLs.

To obtain sets of tissue-specific eQTLs, we first took the union of all eQTLs for tissues in set *T*, given by:

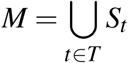

where *S*_*t*_ is the set of eQTLs in tissue *t*. We defined the set of tissue-specific eQTLs for each tissue as the list of significant eQTLs that were significant in only that tissue.

Enrichment analysis was performed by taking the set of signals assigned to each tissue *t* ∈ *T* at each stringency threshold. Each tissue-assigned signal (i.e. 99% genetic credible set) was then mapped to the corresponding GWAS index SNP reported in Mahajan et al. 2018b., yielding a set of index SNPs for each tissue *t*. The program SNPsnap (Broad Institute, accessed Aug 21, 2019) was used to generate 1,000 matched sets of SNPs from the European (EUR) Phase 3 reference panel from the 1000 Genomes Project^23^ using the following parameters: MAF maximum deviation of 5%; maximum gene density deviation of 20%; distance to the nearest gene maximum deviation of 20%; and maximum deviation for the number of LD proxies (i.e. LD “buddies”) set to 20% with a LD threshold of 0.5.

For each tissue *t*, fold enrichments were estimated by taking the observed number of tissue-specific eQTLs among the set of tissue-assigned signals for tissue *t* divided by the mean number of overlapping signals across the 1,000 permuted sets of matched SNPs corresponding to the set of signals (i.e. mapped index SNPs) assigned to tissue *t*. Empirical p-values were calculated by:

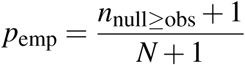

where *n*_null≥observed_ is the number of instances where the number of overlapping tissue-specific eQTLs among a null set of matched SNPs was greater than or equal to the number observed among the set of tissue-assigned signals and *N* is the total number of permutations.

### Functional fine-mapping

A set of comparative *functional* fine-mapping analyses were performed using the program fgwas (v 0.3.6) and the summary statistics from the GWAS meta-analysis for T2D unadjusted for BMI^3^ and three annotation schemes:

- *null* analysis without any genomic annotations
- *multi-tissue* combined analysis using 13-state chromatin state maps for islet, liver, skeletal muscle, and subcutaneous adipose tissue from Varshney et al. 2017^19^ (described above in section “Partitioning chromatin states”).
- *deep islet* analysis based on 15-state chromatin segmentation map for human islet from Thurner et al. 2018^24^; notably, these states were based on a richer set of input features assayed in islets that included ATAC-seq and whole-genome bisulfite sequencing, in addition to histone ChIP-seq.

For both the *multi-tissue* and *deep islet* analysis, fgwas was used to obtain a ‘full model’ by first seeding a model with the single annotation that yielded the greatest model likelihood in a single annotation analysis. This model was extended by iteratively adding annotations - in descending order based on their model likelihoods - until the incorporation of additional annotations no longer increased the model likelihood of the joint model. The ‘full’ model resulting from this procedure was then reduced by iteratively dropping annotations that yielded an increased cross-validated likelihood upon their exclusion from the joint model. The “best joint model” was obtained when this process no longer improved the cross-validated likelihood. The annotations remaining in the “best joint model” were then carried forward for functional fine-mapping.

In the next step, a locus partitioned analysis was performed using the set of annotations from the “best joint model” for the *multi-tissue* and *deep islet* analysis, or no annotations for the null analysis. The default behaviour of fgwas involves partitioning the genome into ‘blocks’ of 5,000 SNPs and assuming no more than one causal variant per block. To account for allelic heterogeneity at loci with conditionally independent signals and to facilitate a comparison with the 99% genetic credible sets (that were constructed using conditionally deconvoluted credible sets), the genome was partitioned into 1 Mb windows centered about each index variant (specified using the –bed command) and fgwas was run using the appropriate set of input annotations for each of the three analytic schemes. Windows involving multiple independent signals required separate fgwas runs, each corresponding to the appropriate set of approximate conditioned summary statistics (i.e. conditioning on the effect of one or more additional signals at a locus)^3^. The resulting PPA values for each SNP in each partitioned ‘block’ was used to construct 99% functional credible sets by ranking SNP by PPA in descending order and retaining those that yielded a cumulative PPA ≥ 0.99.

To compare the differences in fine-mapping resolution between the *multi-tissue* and *deep islet* schemes, at each signal, the difference between maximum 99% functional credible set PPA for each scheme with that resulting from the *null* analysis was obtained as a baseline. These differentials over the null were then compared between the multi-tissue and *deep islet* schemes and significance was assessed using the Wilcoxon rank-sum test. Comparative tests were performed for each set of tissue-assigned signals across the four stringency thresholds.

### Gene co-expression

Genes with TPM counts < 0.1 in > 50% of samples per tissue were excluded and the remaining genes were ranked based on their mean expression across all tissues. For each set of tissue-assigned genetic signals, at each specified classifier threshold, a set of genes was determined based on nearest proximity to the index SNP for each signal. Signals that corresponded to 99% genetic credible sets where coding variants accounted for a cumulative PPA ≥ 0.1 were excluded from the analysis. A *background* set of genes was then obtained by including all genes with rank values +/- 150 about the rank values of each gene in the filtered set. Null sets of genes were then delineated by sampling genes from the *background* set that had rank values within 100 of those for each gene in the gene set. This last step was repeated to generate 1,000 sets of null genes. To assess coexpression in each of the 54 tissues, the rank sum of the genes in the set was recorded and compared with the mean rank sum across the 1,000 sets of null genes separately for each tissue. An empirical p-value was determined with the equation:

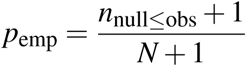

where *n*_null≤obs_ is the number of instances when the rank sum of genes in a null set was less than or equal to the observed rank sum in a given tissue and *N* is the number of permutations. To gauge the magnitude of coexpression, an enrichment factor was defined by taking the mean rank sum across the null sets divided by the observed rank sum. This procedure was repeated for sets of the second and third nearest genes to each index SNP corresponding to tissue-assigned signals across classifier thresholds.

### Physiological cluster enrichment

A set of T2D-associated SNPs that were clustered into physiology groups were obtained from a recent study^25^. As previously described, summary statistics (Z-scores) for a range T2D-relevant metabolic traits (e.g. anthropometric, lipid, and glycemic) were used to cluster 94 coding and non-coding SNPs associated with T2D using “fuzzy” C-means clustering of Euclidean measures^25^. An additional, and partially overlapping, set of 94 T2D-associated SNPs was also accessed and was previously clustered into physiology groups using an input set of sample size-adjusted Z-scores corresponding to 47 T2D-related traits and nonnegative matrix factorization (bNMF) clustering^26^. As not all of the physiologically-clustered SNPs were present among the set of index SNPs corresponding to the 380 fine-mapped genetic association signals, pairwise LD was measured between all SNPs in these sets using the LDproxy tool on the LD Link website (https://ldlink.nci.nih.gov/) and all European populations from the 1000 Genomes Project (Phase 3) as a reference. Physiologically-clustered SNPs were assigned to fine-mapping index SNPs based on maximum pairwise LD where *r*^2^ *>* 0.3. From this approach, 82/94 SNPs and 63/94 SNPs from the two sets of physiologically-clustered signals (from Mahajan et al. 2018a. and Udler et al. 2018., respectively) were mapped to fine-mapped signals in Mahajan et al. 2018b. For each set of tissue-assigned signals with *n* signals, assigned at each classifier threshold, null SNP sets were generated by randomly sampling *n* signals from the set of 380 fine-mapped signals. A null distribution was obtained by generating 10,000 null sets and recording the overlap of null signals with each of the physiologically-clustered signals. An empirical p-value was obtained with the equation:

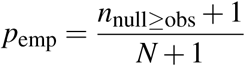

Where *n*_null≥obs_ is the number of instances where the observed overlap between a null set and a reference set of physiologically assigned signals was greater than or equal to the observed value for the query set of tissue-assigned signals and *N* is the total number of null sets (i.e. 10,000). An enrichment factor was obtained by taking the observed overlap divided by the mean of the null overlap values.

### Enrichment for trait-associated SNPs from GWAS

GWAS summary statistics for all available traits and diseases were downloaded from the NHGRI-EBI GWAS catalogue (https://www.ebi.ac.uk/gwas/; v1.0; accessed Aug 23, 2019). Coordinates for all trait-associated SNPs in the catalogue were mapped to genome build GRCh38. GRCh38 coordinates for index SNPs corresponding to each of the 99% genetic credible sets were obtained from the Ensembl website (https://www.ensembl.org/) by querying with reference SNP id number. Proxy SNPs were determined for each SNP in the set of index SNPs corresponding to the 99% genetic credible sets by using the –show-tags function in PLINK (v 1.90b3) to identify SNP proxies with linkage disequilibrium (LD) *r*^2^ ≥ 0.8 among a reference panel of European individuals from the 1000 Genomes Project (Phase 3). VCF files for SNPs from the 1000 Genomes Project mapped to genome build GRCh38 were downloaded from the project website (http://ftp.1000genomes.ebi.ac.uk/). For each set of tissue-assigned signals, enrichment was assessed across each of the 3,616 diseases or traits in the GWAS catalogue. The observed number of SNPs overlapping the set of index and proxy SNPs corresponding to the tissue-assigned signals and the set of trait-associated SNPs for a given GWAS was recorded. To obviate bias due to local LD, multiple SNPs (i.e. index and proxies) corresponding to a single signal that were shared with the set of GWAS SNPs were recorded as a single overlap for that signal. A null distribution of SNP overlaps was obtained through 10,000 rounds of random sampling from the set of index SNPs corresponding to each of the 380 fine-mapped credible sets. An empirical p-value was obtained with the formula:

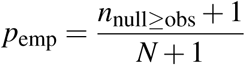

where *n*_null≥obs_ is the number of instances where the number of SNP overlaps between a null and GWAS SNP set exceeded the observed overlap for the set of tissue-assigned signals. The magnitude of enrichment was measured by the number of observed overlaps divided by the mean of the overlaps across the null sets.

## Results

### An integrative approach for obtaining *tissue-of-action* scores at trait-associated loci

We set out to quantify, in the form of TOA scores, the contribution of disease-relevant tissues to each genetic association signal from a recent GWAS meta-analysis of T2D by integrating genetic, genomic and transcriptomic data. To do this, we developed a scheme that derived, for each GWAS signal, a measure of overlap with tissue-specific regulatory annotations, and then combined these, using weights derived from both genetic fine-mapping and genome-wide measures of tissue- and annotation-specific enrichment (**Figure 1**).

We used chromatin states from a recent study^19^ to form a reference set of epigenomic annotations focusing on tissues involved in insulin secretion (pancreatic islets) and insulin-response (skeletal muscle, subcutaneous adipose, and liver) that play central roles in the pathophysiology of T2D. There is support for the role of these tissues from patterns of overall genome-wide enrichment of tissue-specific regulatory features and from the known effects at the subset of T2D association signals for which causal mechanisms have been established^13, 14, 19, 24, 27^.

To obtain tissue scores at each genetic signal, we first delineated a set of annotation vectors based on the physical position of each SNP in the corresponding 99% genetic credible set (from Bayesian fine-mapping) with respect to the panel of tissue-specific chromatin states (**Figure 1**). For non-coding SNPs, binary values were used to encode genome mapping (i.e. whether or not a SNP maps to a regulatory region in a given tissue as shown in Step 1A in **Figure 1**). For the minority of credible set SNPs that map to coding sequence, quantification focused on measures of tissue-specific RNA expression for the genes concerned to further inform the relative importance of the evaluated tissues (see Methods) (**Figure 1**, Step 1B).

Next, we combined and scaled the annotation vectors to yield a vector of *tissue* scores that were used to partition the PPA of each credible SNP (Step 2). To facilitate this partitioning and to account for the relative importance of relevant tissues with respect to overall T2D pathogenesis, we first estimated genome-wide enrichment of T2D-associated SNPs across a set of tissue-specific genomic annotations. We used the enrichment values as weights to adjust the relative tissue contributions of SNPs mapping to distinct functional annotations or to functional annotations shared in more than one tissue (see Methods) (**Supplementary Figure 1A-C**). This allowed us, for example, to upweight the islet contribution, relative to that for skeletal muscle, for SNPs mapping to enhancers shared between these tissues to account for the different genome-wide enrichment priors observed for these tissues.

**Figure 1.**
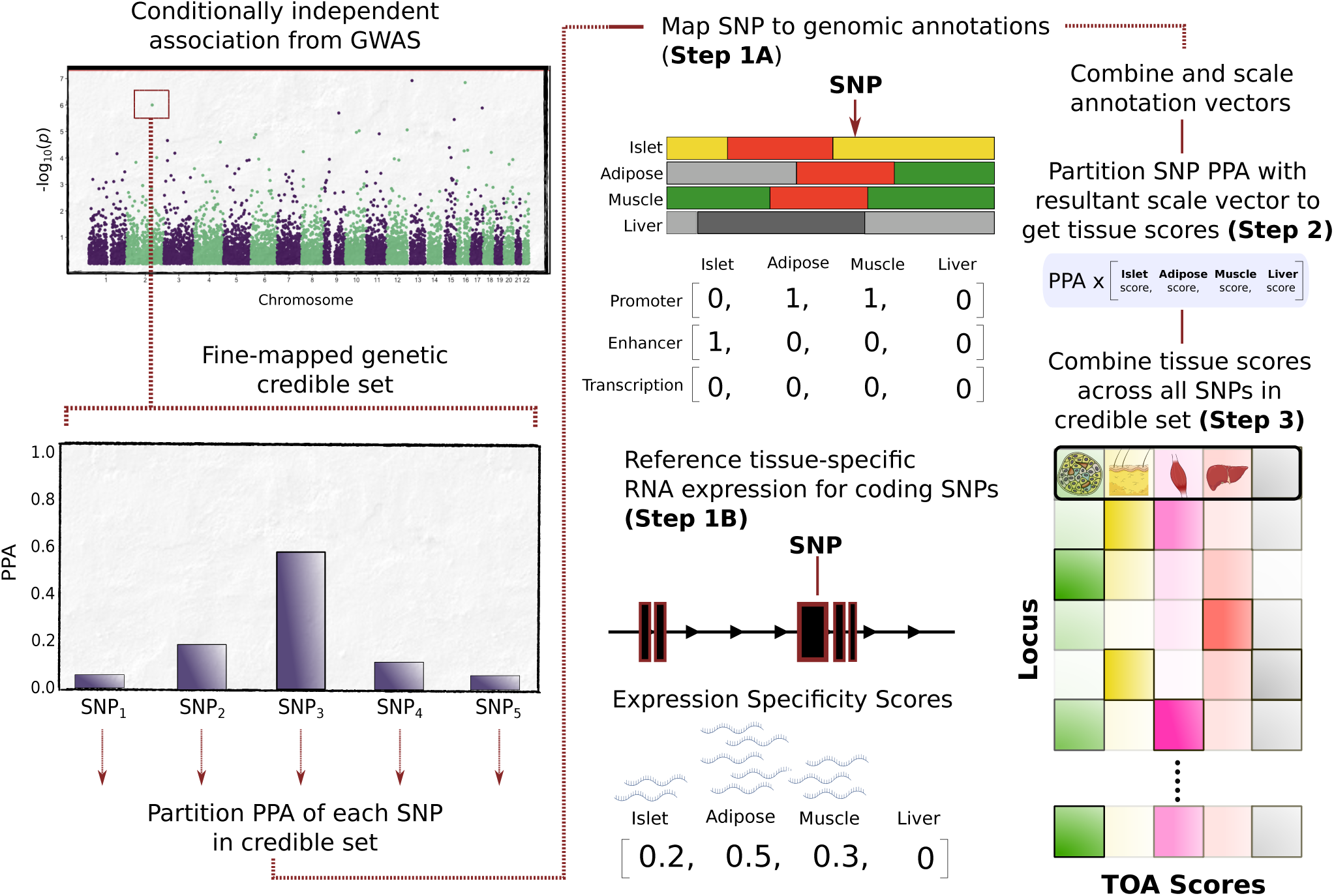
Systematic approach for obtaining tissue-of-action scores. Fine-mapping of conditionally-independent GWAS signals results in a set of credible variants, each with a posterior probability of association (PPA). The illustrated example shows a signal with five SNPs in its credible set with SNP_3_ as the variant with the maximum PPA. Each credible SNP is then mapped to a panel of chromatin state annotations across four disease-relevant tissues to obtain a set of annotation vectors (**Step 1A**). An additional annotation vector for SNPs mapping to coding sequence (CDS) is obtained from expression specificity scores (ESS) calculated from gene expression levels across the four tissues (**Step 1B**). The set of annotation vectors for each SNP are then summed and scaled, yielding a vector used to partition the PPA value (**Step 2**). The resultant vectors for each SNP in a genetic credible set are then summed and scaled to yield a tissue-of-action (TOA) score for each tissue at the GWAS signal corresponding to the credible set (**Step 3**). Any residual PPA values from SNPs not mapping to any of the evaluated tissue annotations are allocated to an “unclassified” score (grey column in matrix).

Across all tissues, we found that the active transcription start site (TSS) annotation, distinguished by strong ChIP-seq signal for H3K27ac and H3K4me1 histone modifications, was the most consistently enriched feature (log2 fold enrichment from 2.46 to 2.79) (**Supplementary Figure 1A-B**). However, the most highly-enriched single annotation detected involved type 1 active enhancers in human islets (as characterized by H3K27ac and H3K4me3) (log2 FE=2.84, 95% CI, 1.48-3.62). Coding sequence was also highly enriched for T2D-associated variants (log2 FE=2.59, 95% CI, 2.08-3.01) (**Supplementary Figure 1B**).

In the final step, the tissue partitioned PPA values were combined across all SNPs in the credible set to yield a set of TOA scores for each association signal which preserves the information captured by the fine mapping (**Figure 1**: Step 3). PPA values corresponding to SNPs not mapping to active regulatory annotations in any of the four evaluated tissues (e.g. repressed or quiescent regions) were allocated to an “unclassified” score (see Methods). The resulting set of TOA scores for each genetic signal captures the strength of genetic, genomic, and transcriptomic evidence that the signal acts through each of the evaluated tissues. Using this framework, we calculated TOA scores for each of the 380 fine-mapped T2D signals (**Supplementary Table 1**).

### *Tissue-of-action* scores support a key role for strong enhancers in human islets

By combining TOA scores across all 380 signals, we estimated the relative contribution of each tissue to the overall genetic risk of T2D reflected across fine-mapped loci. Islet accounted for the largest share of the cumulative TOA score (29%) with markedly lower contributions from liver, adipose, and skeletal muscle (**Figure 2A**, inset). Across the 380 loci, 80% of the cumulative TOA score was attributable to SNPs mapping to coding regions or to active chromatin states in these four tissues (**Figure 2A**). Within this fraction, SNPs mapping to weakly transcribed regions accounted for the largest share (51%) relative to those mapping to coding and other regulatory annotations (**Figure 2A**). Overall, weakly transcribed regions account for 23% of the genome (ranging from 22% in skeletal muscle to 26% in islet), and are generally located near other more active annotations (**Supplementary Figure 1D**).

**Figure 2.**
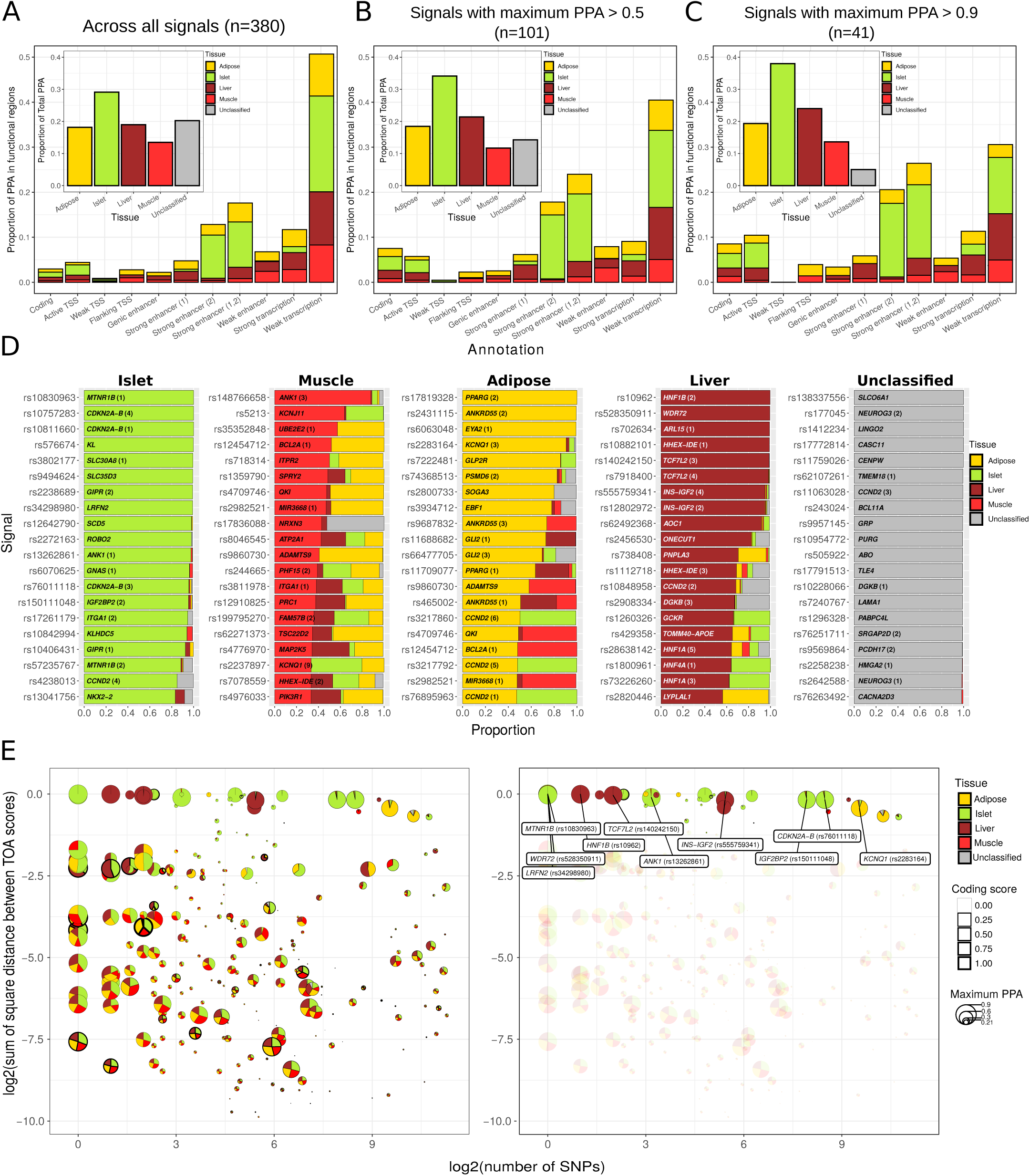
The profile of tissue-of-action scores across T2D signals. **A)** The proportion of total PPA summed across all 380 signals is shown for each tissue (inset). The proportion of total PPA is also shown for each annotation group (outset). Proportions are also exhibited for the subset of signals with maximum credible set PPA > 0.5 in panel **B** and for the subset of signals with maximum PPA > 0.9 in panel **C. D)** The profile of TOA scores is shown for the top 20 signals ranked for each tissue. The locus name and rs accession number for the index SNP is indicated for each signal. Signals at loci with multiple conditionally-independent signals are indicated by parenthetical numbers (i.e. one is primary signal, two is secondary signal, etc.). **E)** Relationship between fine-mapping resolution and TOA score diversity. Log2 of the number of credible SNPs for each fine-mapped signal is shown on the x-axis and the log2 value of the sum of square differences between TOA scores for each signal is shown on the y-axis (i.e. higher values on the y-axis correspond to greater tissue “specificity”). The profile of TOA scores are indicated within pie charts where the diameter of each circle corresponds to the maximum PPA for the credible set. The line thickness for each circle indicates a coding score for each credible set (i.e. the proportion of cumulative PPA attributable to coding variants). The left panel shows all credible sets with unclassified scores < 0.10 (n=259) and the right panel highlights the subset of “tissue-specific” signals with TOA scores ≥ 0.8. The ten “tissue-specific” signals with the highest maximum credible set PPA are labeled in the right panel.

Crucially, credible sets vary markedly in their fine-mapping resolution (median credible set size 42 SNPs, range 3997 SNPs: median maximum PPA value 0.24, range 0.01-1.0). We reasoned that the estimates for weakly transcribed regions (and for annotations to tissues outside the four most relevant to diabetes) were likely inflated by incomplete fine-mapping: less resolved credible sets involving multiple SNPs are likely to map to disparate annotations across tissues. When we evaluated the 101 signals with maximum PPA>0.5, the TOA score proportions attributed to weak transcription and unclassified proportions decreased to 40% and 14%, respectively (**Figure 2B**). These proportions further decreased amongst the 41 signals with maximum PPA>0.9 (31% and 5% respectively) (**Figure 2C**). In contrast, the relative contribution of SNPs mapping to strong enhancers increased with greater fine-mapping resolution (from 18% to 26%) (**Figure 2A-C**). In particular, the contribution for strong enhancers in islet was disproportionately high among the most finely-mapped signals and underscores a prominent role for these regulatory regions in T2D risk (**Figure 2C**).

Although the relative TOA score proportions varied with fine-mapping resolution, the contribution from islet was consistently greater than that for liver, adipose, or muscle (by a factor of 1.5) (**Figure 2A-C**, inset). Notably, for credible SNP mapping to strong enhancers, the relative TOA proportions were considerably higher for islets (57-63%) than for adipose (18-24%), liver (14%), and skeletal muscle (5-6%). Increasing fine-mapping resolution tracked with increasing evidence that causal variants were disproportionately concentrated in islet strong enhancers (**Figure 2A-C**, outset). When we additionally weighted TOA scores by the adjusted GWAS effect size for each signal (see Methods), the overall islet contribution increased further, albeit slightly, from 29% to 31% across all signals (**Supplementary Figure 2D-F**). Overall, the profile of tissue-of-action scores (particularly across more signals with greater fine-mapping resolution) recapitulates the epigenomic architecture of T2D derived from earlier studies, which have indicated that regulatory annotations in islets - and strong enhancers in particular - are particularly important (**Figure 2B-C**).

### Distinct TOA profiles indicate pleiotropic effects in multiple tissues

The prime motivation for generating TOA scores was to identify the tissues that most likely mediate disease risk at each genetic signal. We first sought to identify signals where only a single tissue was likely relevant to disease risk. We found that 10% (39/380) signals had profiles where the TOA score for one of the four tissues exceeded a threshold of 0.8, consistent with predominant action in a single tissue: 21 of these involved primary or unique signals at their respective loci whereas the remaining 18 arose from secondary signals at loci with multiple independent signals (Supplementary Table 1). Among the primary signals, 14 mapped to islet (including signals at *MTNR1B, SLC30A8, CDKN2A/B* loci), five to liver (e.g. *AOC1, WDR72*), and two to adipose (*EYA2, GLP2R*) (**Figure 2D-E**). No primary signal met this criterion for skeletal muscle: the signal with the highest TOA score for skeletal muscle (0.88) corresponded to a secondary signal (rs148766658) at the *ANK1* locus (**Figure 2D**). The proportion of signals with TOA profiles consistent with a single tissue of action increased with greater fine-mapping resolution (17/101 or 16% of signals with maximum PPA≥0.5) (**Supplementary Table 1**).

Aside from these 39 signals, calculated TOA scores for most T2D signals revealed substantial contributions from multiple tissues. We reasoned that this apparent “tissue sharing” could have arisen for two main reasons. The first involves a highly resolved signal from genetic fine-mapping at which the causal variant maps to a single regulatory element active in multiple tissues. The second occurs when a lower resolution signal encompasses many credible set variants that map to distinct regulatory elements with different patterns of tissue specificity. There was some evidence in favor of the latter: maximum credible set PPA values positively correlated with the SSD between TOA scores (i.e. more refined credible sets corresponded to higher measures of tissue specificity) (Adj.*R*^2^=0.04, p-value=9.8*x*10^−5^, **Supplementary Figure 3**). However, the magnitude of the effect of fine-mapping resolution on tissue specificity was small (the beta coefficient for the regression of SSD on maximum PPA was 0.17). We conclude that differences in fine-mapping resolution alone do not account for the extent of “tissue-sharing” observed across T2D signals, implying that many signals involved regulatory elements shared across tissues.

**Figure 3.**
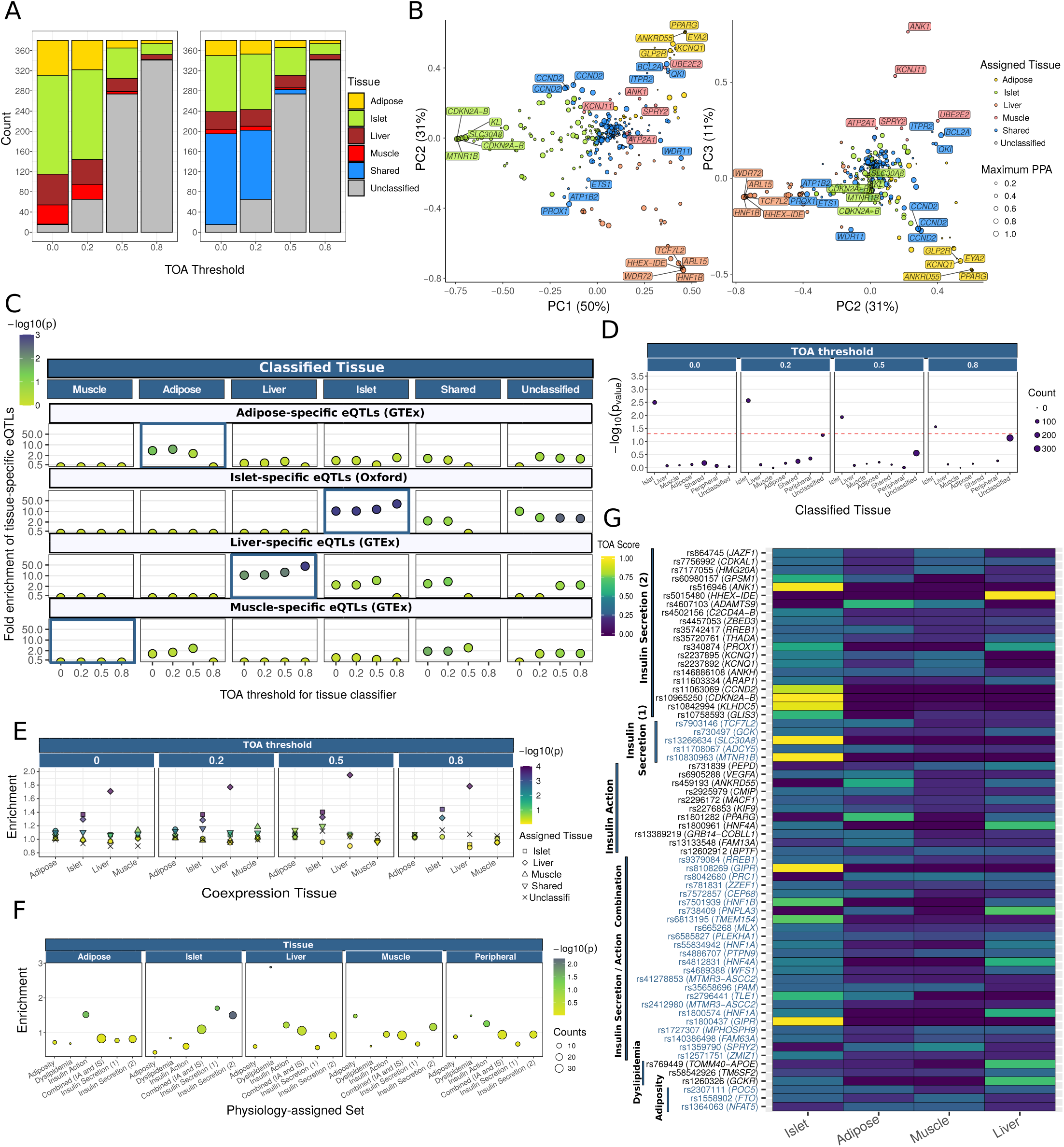
Enrichment of tissue-specific epigenomic and physiological features among classified signals. **A)** Number of signals assigned to each tissue by the classifier for each of the four TOA score thresholds: 0.0, 0.2, 0.5, and 0.8 (left panel). Signal counts are shown across thresholds using a classifier that assigns signals with two or more TOA scores within 0.1 of each other as “shared” signals (right panel). **B)** PCA plots of the decomposition of the TOA score matrix comprising the 306 signals with “unclassified” scores ≤ 0.5. Each point corresponds to a signal where the size indicates the maximum credible set PPA and the color indicates the assigned tissue at the TOA score threshold ≥ 0.2 using the classifier that included a “shared” designation. **C)** Selective enrichment of tissue-specific eQTLs among credible sets for signals assigned to subcutaneous adipose, islet, liver, and skeletal muscle tissue. Color indicates significance of enrichment. **D)** Selective improvement in fine-mapping resolution at islet-assigned signals when richer islet chromatin states are deployed. Comparison of *functional* fine-mapping resolution using a panel of chromatin state annotations based on histone ChIP-seq across the four T2D relevant tissues versus chromatin states based on islet ChIP-seq, ATAC-seq, and DNA methylation (WGBS). **E)** Coexpression of nearest genes annotated to sets of tissue-assigned signals across stringency thresholds. Shape indicates the tissue to which the set of signals were assigned. **F)** Selective TOA score enrichment within relevant sets of physiology-assigned signals. Size corresponds to the number assigned signals in each physiology group. **G)** Tile plot of TOA scores for physiologically-assigned signals. Signals are ordered by physiology group and the corresponding GWAS locus is shown.

To explore this further, we considered signals likely to involve shared effects across tissues on the basis that the difference between the two highest TOA scores was <0.10 (**Supplementary Table 2**). The resulting set of “shared” signals conspicuously spanned the range of mapping resolution, as indicated by the number of credible SNPs and maximum PPA for each signal (**Figure 2E**). There were eight signals that were fine-mapped to a single credible SNP (i.e. maximum PPA>0.99) and most clearly demonstrated tissue-shared regulation. This included the primary, non-coding signal at the *PROX1* locus (rs340874) with effects in both islet (TOA=0.50) and liver (TOA=0.49): the index SNP at this signal (PPA=1.0) mapped to a common active transcription start site in these tissues (**Supplementary Figure 4A, Supplementary Table 2**). This set also included primary signals at the *RREB1* (rs9379084; islet TOA=0.31; adipose TOA=0.27; muscle TOA=0.22), *CCND2* (rs76895963; islet TOA=0.53; adipose TOA=0.47), and *BCL2A* (rs12454712; muscle TOA=0.52; adipose TOA=0.48) loci (**Supplementary Figure 4A, Supplementary Table 2**). There were an additional 33 signals with apparent tissue-sharing where the fine-mapping resolution was somewhat less precise (maximum PPA>=0.5). These included the primary signal at the *TCF7L2* locus (rs7903146; adipose TOA=0.37; islet TOA=0.31) and secondary signals at *HNF4A* (rs191830490 [liver TOA=0.40, islet TOA=0.31] and rs76811102 [islet TOA=0.32, muscle TOA=0.25, liver TOA=0.24]) (**Supplementary Figure 4B, Supplementary Table 2**). Amongst the total of 101 signals at which the fine-mapping resolution was such as to identify a lead SNP with PPA exceeding 0.5, 41% had evidence that they might involve regulatory effects in two or more tissues.

**Figure 4.**
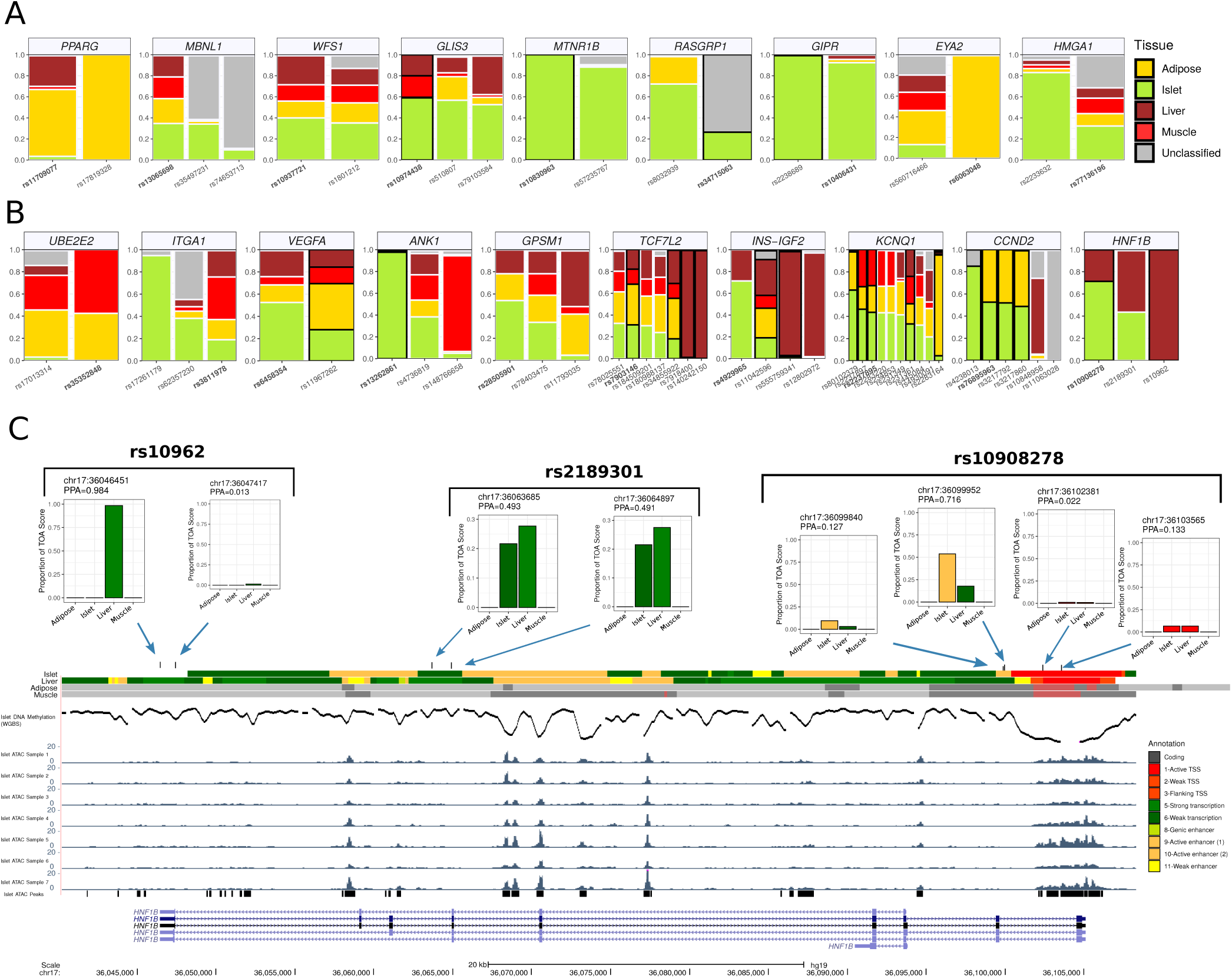
Multiple tissues implicated by epigenomic scores at heterogenous loci. **A)** Profile of TOA scores for the nine loci with all signals receiving identical, “non-shared” tissue assignments at the 0.2 stringency threshold **B)** Profile of TOA scores for the ten loci with all signals receiving distinct, “non-shared” tissue assignments at the 0.2 threshold. **C)** epigenomic profile of PPA values attributable to each credible SNP of the primary signal at the *HNF1B* locus. For each credible SNP, the PPA value attributable to each tissue annotation is shown along with its position on chromosome 17 (genome build hg19). Chromatin state maps for islet, adipose, muscle, and liver tissue from Varshney et al. 2017. are shown along with ATAC-seq tracks for seven representative islet samples, called ATAC-seq peaks from a set of islet ATAC samples (n=17), and DNA methylation (whole genome bisulfite sequencing) in human islets from Thurner et al. 2018.

### A rule-based classifier for assigning fine-mapped signals to tissues

As *tissue-of-action* scores appeared to distinguish specific from shared signals (**Figure 2D-E**), we implemented a rule-based classifier that assigns signals to tissues according to their TOA scores across a range of stringencies. A GWAS signal was assigned to a tissue if that tissue had the highest TOA score and exceeded a specified TOA threshold (ranging from permissive thresholds of zero and 0.2 to more stringent thresholds of 0.5 and 0.8). Consistent with the observation that islet accounted for most of the cumulative PPA across loci (**Figure 2A-C**), more signals were assigned to islet than to liver, muscle, or adipose tissue across all TOA thresholds. For example, at a TOA threshold of 0.2, 178 signals (47%) were classified as islet whereas a total of 137 signals (36%) were assigned to insulin-responsive peripheral tissues (58 adipose, 49 liver, 30 muscle) (**Figure 3A**, left panel). Given the extent of tissue sharing observed across signals, we adapted the classifier scheme to allow for a shared category (defined as above): at the same TOA threshold, this yielded 110 islet, 33 liver, 27 adipose and 8 muscle signals, plus 137 shared signals (**Figure 3A**, right panel). These proportional differences between islet, muscle, adipose, and liver were maintained across TOA thresholds (**Figure 2D**). For example, the distribution of the 39 signals classified at the 0.8 threshold included 22, 10, 6 and 1 signals classified as islet, liver, adipose, and muscle, respectively (**Figure 2D**).

Principal component analysis of these data revealed that most variation in TOA scores (50%) distinguished islet signals from those assigned by the classifier to insulin-responsive peripheral tissues, consistent with the distinct functions of these tissues in regulating glucose homeostasis (**Figure 3B**). The distinction between liver and adipose signals accounted for a further 31% of variation. Signals classified as shared mapped between the clusters of tissue-assigned signals (**Figure 3B**). For example, three of the six conditionally-independent signals at the *CCND2* locus (including the primary signal at rs76895963; PPA=1.0) classified as “shared”, and mapped equidistant between adipose and islet clusters (**Figure 3B, Supplementary Table 1**). Other clear examples include the primary signals at the *PROX1* and *BCL2A* loci described above that exhibit profiles with sharing between islet and liver, and muscle and adipose, respectively (**Figure 3B**).

Despite incorporating data from the four tissues most relevant to T2D pathogenesis, a considerable number of signals remained unclassified across stringency thresholds (e.g. 65 signals at the 0.2 threshold), reflecting the appreciable proportion of cumulative PPA at these signals attributable to credible set SNPs that did not map to active regulatory regions in any of these tissues. This can, in part, be explained by the poorer fine-map resolution of these signals compared to classified signals (median credible set size: 57 versus 36 SNPs; median maximum PPA: 0.20 vs. 0.25). However, it is possible that some of the unclassified signals involve tissues or cell types not explicitly included in our analysis. Indeed, signals that remained unclassified at the TOA score ≥0.2 threshold were more likely to map to regions that were actively repressed or quiescent (i.e. low signal) in the four evaluated tissues (**Supplementary Table 3**).

Given that a subset of T2D signals are driven by adiposity and presumed to act through central mechanisms^3^, one obvious omission from the tissues considered in our primary analysis was brain (or, more specifically, hypothalamus). For example, T2D-associated variants at the obesity-associated *MC4R* locus (encoding the melanocortin 4 receptor) were assigned as unclassified in our analyses^3, 28–31^. However, using chromatin state maps from multiple brain regions, we found a deficit, rather than an excess, of PPA enrichment amongst active enhancers (0.032 vs.0.147; p-value=7.5*x*10^−5^) and promoters (0.007 vs.0.043; p-value=0.0054) for unclassified signals (as compared to classified) (**Supplementary Table 3**). The data available did not, however, include chromatin state maps for the hypothalamus. Overall, it is to be expected that classification of currently-unclassified signals will improve with increased fine-mapping resolution and the availability of detailed chromatin annotations from additional tissue and cell types.

### Tissue-assigned signals are validated by orthogonal tissue-specific features

We sought to validate the performance of the classifier by evaluating how assignments from the TOA classifier matched tissue-specific information from three orthogonal sources: tissue-specific eQTL enrichment, “functional” fine-mapping, and proximity-based gene coexpression analysis of non-coding signals. For these evaluations, we used the version of the classifier that allows for a shared designation.

To determine if tissue-assigned signals were matched to tissue-specific eQTLs, we assembled *cis*-eQTLs for liver, skeletal muscle, subcutaneous adipose tissue (all GTEx V7) and human islets^5^, and defined sets of tissue-specific eQTLs (see Methods). The set of signals assigned by the TOA classifier to islets were significantly, and selectively, enriched for islet-specific eQTLs across all TOA thresholds (ranging from 10-fold to 31-fold enrichment [p-values<0.001]) as compared to matched sets of SNPs (see Methods) (**Figure 3C**). Similarly, the set of signals assigned by the TOA classifier to liver showed marked, selective, enrichment for liver-specific eQTLs across TOA thresholds (**Figure 3C**). Overall, the more confidently assigned genetic signals retained at more stringent TOA thresholds tended to have larger point effect estimates, though the reduced number of signals meeting the more stringent thresholds led to wider confidence intervals and some reduction in the statistical significance of the enrichments. Relatively few signals were assigned to adipose and skeletal muscle at higher thresholds (**Figure 3A**): nonetheless, adipose-assigned signals were the most enriched for adipose-specific eQTLs at lower stringency (e.g. 5-fold enrichment, p-value = 0.021, at the 0.2 threshold (**Figure 3C**). In contrast, although sets of signals classified as shared showed some enrichment for tissue-specific eQTLs at less stringent thresholds, these enrichments were generally lower than those for signals assigned to the corresponding tissues (**Figure 3C**). These data indicate that the tissue assignments made by the classifier are consistent with the information from *cis*-eQTL analyses in corresponding tissues.

The second validation analysis was motivated by the use of high-resolution epigenomic maps to improve genetic fine-mapping. For the present study, we had derived TOA scores using chromatin states based solely on ChIP-seq data^19^: this was a conscious decision designed to minimize technical differences in the depth of annotation available between tissues given that chromatin accessibility and DNA methylation data were not as widely available. However, we had previously shown that islet enhancer chromatin states obtained from a segmentation analysis that incorporated information from DNA methylation, ATAC-seq, and histone ChIP-seq data yielded higher enrichment of T2D-associated SNPs than enhancer states delineated from ChIP-seq data alone^24^. We reasoned that accurate assignment of islet signals by the TOA classifier would be expected to result in an improvement in fine-mapping, following the use of fine-grained islet functional information, which was restricted to the set of islet-assigned signals. To test this hypothesis, we performed a comparative “functional” fine-mapping analysis (see Methods) using this richer set of islet annotations^24^ and found that the mean maximum credible set PPA significantly increased for islet-assigned signals relative to the corresponding value from a joint analysis based on ChIP-seq data alone (e.g. mean PPA increase=0.064; p-value=0.0027 at the 0.2 threshold) (**Figure 3D**).

This was true across all TOA thresholds. In contrast, credible sets for signals assigned to insulin-responsive peripheral tissues showed no improvement in fine-mapping resolution with the richer islet annotations (**Figure 3D**). These data indicate that the tissue assignments made by the TOA classifier are consistent with the information from more detailed functional annotations in relevant tissues.

The third validation approach involved assessing genes for overlapping coexpression^32^. Although the genes lying closest to the lead regulatory variants at GWAS signals are not guaranteed to be the causal transcript, the set of “nearest genes” is, nonetheless, likely to be enriched for the genes responsible for mediating such associations^33^. As such, we reasoned that performance of the classifier would be reflected in the extent to which genes near non-coding signals were coexpressed in the corresponding tissue as compared to more distal genes. We assigned a single (nearest) gene to each tissue-classified signal and found that the set of genes nearest to islet-assigned signals showed the most pronounced coexpression in human islet tissue across all TOA thresholds (e.g. p-value=0.0003 at threshold 0.8) (**Figure 3E**) and across an expanded set of tissues, including 53 tissues from the GTEx Project (**Supplementary Figure 5A**). This coexpression signal was lost for the sets of second- and third-nearest genes (**Supplementary Figure 5B-C**). Similar results were observed for liver, muscle and adipose (**Figure 3E**). In contrast to the sets of nearest genes annotated to signals assigned to specific tissues, gene sets annotated to signals classified as either “shared” or “unclassified” did not show pronounced co-expression in any of the evaluated tissues (**Figure 3E, Supplementary Figure 5A**). These data indicate that the tissue assignments made by the classifier are consistent with the information from co-expression analyses in corresponding tissues. Collectively, the data from these three analyses further supports the validity of the *tissue-of-action* scores generated by our approach.

### Tissue-assigned signals are supported by physiological clustering

It is possible to assign T2D risk alleles with respect to physiological impact based on patterns of genetic association with related quantitative traits such as fasting glucose and insulin levels, circulating lipid levels, and anthropometric traits^2, 25, 26, 34, 35^. At the same time, those same physiological processes map to specific tissues (e.g. insulin secretion from pancreatic islets). We asked therefore if the tissue assignment of signals by the TOA-classifier (based on tissue-specific molecular data) was consistent with the assignments made on the basis of whole body physiology. We focused on a set of 82 T2D-associated variants that had previously been partitioned using a “fuzzy” clustering algorithm^25^ to six physiological clusters and were in linkage disequilibrium with lead variants from the set of 380 fine-mapped credible sets (see Methods).

We first asked if these signals assigned to these six physiological clusters differed with respect to their TOA score distributions. Variants assigned to the two *insulin secretion* clusters (characterised by associations with reduced fasting glucose and HOMA-B levels but differing with respect to effects on proinsulin and HDL cholesterol levels) had higher islet TOA scores than variants in the other physiological clusters (enrichment = 1.5, 1.7 [p=0.006, 0.03] for the type 2 and type 1 *insulin secretion* cluster, respectively) (**Figure 3F-G**). Variants assigned to the *insulin action* and *dyslipidemia* clusters corresponded to signals with significantly higher adipose (1.5-fold, p=0.034) and liver scores (2.9-fold, p=0.009), respectively (**Figure 3F-G**). Reciprocally, sets of TOA-classifier tissue-assigned signals were significantly enriched for SNPs from relevant physiology sets (**Supplementary Figure 6A**). Similar results were obtained from a different (but overlapping) set of physiological clusters derived using an alternative clustering scheme^26^ (**Supplementary Figure 6B-C**).

These patterns were confirmed by evaluating enrichment across all phenotypes present in the NHGRI-EBI GWAS catalogue. For example, T2D signals assigned to adipose by the TOA-classifier were enriched for variants associated with traits relevant to fat distribution (e.g waist-to-hip ratio adjusted for BMI, 3.5-fold, p-value<0.0001) whereas signals assigned to liver and islet were enriched for SNPs associated with total cholesterol levels (3.3-fold, p-value=0.0011) and acute insulin response (2.3-fold, p-value=0.009), respectively (**Supplementary Figure 7**). Collectively, these results indicate that tissue assignments based on TOA scores derived from molecular data are consistent with inference based on *in vivo* physiology.

### Epigenomic clustering implicates multiple tissues at loci with independent signals

The 380 fine-mapped genetic credible sets map to 239 loci, 84 of which harboured multiple conditionally-independent signals^3^. As disparate signals within the same locus cannot be assumed, purely on the basis of genomic adjacency, to influence disease risk through the same downstream mechanism, we asked how often the classifier assigned independent signals at a locus to different tissues. We focused on the 0.2 threshold as this allowed us to assign signals to each of the four T2D-relevant tissues, whilst still being widely validated by the approaches described above (**Figure 3**). There were 60 loci where at least two signals were assigned to a tissue or designated as “shared” (**Supplementary Figure 8**), but we focused on 19 loci where two or more independent signals received tissue-specific assignments (rather than “shared”). Of these, there were nine loci where constituent signals were given identical tissue assignments. These included *PPARG* and *EYA2* (all signals designated as adipose) and seven others - including *MTNR1B* and *GIPR* – at which all signals were assigned to islet (**Figure 4A**).

This left ten loci where there was divergent assignment of signals. One of the clearest examples involves the *HNF1B* locus where three signals (each comprising non-coding variants) varied markedly in their TOA scores from islet and liver (**Figure 4B**). The lead signal, at rs10908278, was assigned to islet as the credible variants with the highest PPAs (0.72 and 0.13) both mapped to the same strong islet-specific enhancer (**Figure 4C**). In contrast, the rs10962 signal was assigned to liver as the likely causal variant (PPA=0.98) mapped to a strongly transcribed region specific to liver. The remaining signal, at rs2189301, was classified as “shared” as the principal credible set variants (both with PPA=0.49) mapped to a transcribed region in both islet and liver, with the latter showing a stronger epigenomic signature for transcription (**Figure 4C**).

Large-scale GWAS meta-analysis in Europeans has uncovered multiple signals at the *ANK1* locus. One of these, at rs13262681, colocalises with an eQTL for *NKX6*.*3* expression in pancreatic islets^3^. Using the TOA-classifier, we found that this signal (rs13262861; PPA=0.97) was designated as islet given overlap with a strong islet enhancer. On the other hand, an independent signal at rs148766658 (43 Kb from rs13262861) was categorized as a muscle signal as credible set SNPs (maximum PPA=0.25) mapped to strong enhancer and transcribed chromatin states in skeletal muscle (**Supplementary Figure 9A-B**). These data suggest that this “locus” is really a composite of overlapping associations, with entirely distinct effector transcripts and tissues-of-action. Notably, a recent GWAS meta-analysis of T2D in 433,530 East Asians has uncovered independent signals in this region that distinctly colocalize with either an eQTL for *NKX6-3* in islet or an eQTL for *ANK1* expression in skeletal muscle and subcutaneous adipose tissues^36^. Although there is incomplete LD between the specific *ANK1* variants detected in the European and East Asian meta-analyses (between the secondary signals in particular), our results are consistent with the presence of distinct signals near *ANK1* with disparate tissue effects. This example highlights the growing limitations of segmenting the genome into loci, based purely on measures of adjacency, with component signals at each locus considered to share some functional relationship. Instances such as this, where proximal signals represent functionally distinct mechanisms, indicate that such assumptions can be misleading and are likely to become less tenable as the density of GWAS hits for each disease of interest increases.

Amongst the ten loci displaying evidence for “tissue heterogeneity” across signals was *TCF7L2*. Of the seven independent signals at *TCF7L2* revealed by conditional fine-mapping, two (at rs7918400 and rs140242150) were assigned solely to liver (**Figure 4B**). The remaining five signals revealed contributions from both islet and adipose (**Figure 4B**). This group includes the lead signal at *TCF7L2* (lead SNP, rs7903146), which remains the strongest common variant T2D association in Europeans. This signal was classified as “shared”, with similar TOA scores from islet (0.31) and adipose (0.37). Crucially, this signal did not fine-map exclusively to rs7903146 (PPA=0.59; MAF=0.26) in Europeans: the 99% credible set included two additional SNPs3. Variant rs34872471 (PPA=0.36) is in near perfect LD (r2=0.99) with rs7903146 in Europeans^23^. While rs7903146 has a pronounced islet signature due to mapping to an epigenetically active region in islet (a strong enhancer with high chromatin accessibility and low DNA methylation), rs34872471 mapped to a strong enhancer active only in adipose (**Supplementary Figure 9C**). The net effect, based on this information, is a “shared” designation. In truth, either there is a single causal variant at this locus (rs7903146, or potentially, rs34872471) and once resolved, this signal can be correctly assigned to the relevant tissue; or both SNPs are directly contributing to T2D risk through distinct mechanisms in islet and adipose tissue.

### TOA scores advance resolution of effector transcripts

Given the TOA-score classifier was able to discriminate sets of genetic signals that were supported by orthogonal validation features, we next considered the value of TOA scores to clarify regulatory mechanisms and enhance the identification of downstream effector transcripts at T2D-associated loci. One widely used approach for promoting candidate causal genes at GWAS loci involves identifying *cis*-eQTL signals that colocalize with trait-associated SNPs^37, 38^. However, *cis*-eQTL signals show appreciable tissue specificity, raising the possibility of misleading inference if analyses are conducted in a tissue irrelevant to the signal of interest^39, 40^. For example, a *cis*-eQTL specific to liver is likely to be more informative for a T2D signal assigned to liver, than one assigned to islet.

We explored the utility of incorporating TOA scores for T2D-relevant tissues into a previous colocalization analysis^3^ fine-mapping resolution, we evaluated eQTL colocalisation results involving the 101 T2D GWAS signals with credible sets featuring lead SNPs with maximum PPA>=0.5. A total of 378 eQTL colocalizations (eCaviar CLPP*>*=0.01) were detected across 53 signals with a median of four colocalizations (implicating four distinct pairs of tissues and eGenes) per signal (**Supplementary Table 4**). At some loci, the number of colocalizations detected can be substantial: at the *CLUAP* locus, for example, the lead T2D SNP (rs3751837, PPA=0.90) was the source of 64 cis-eQTL colocalizations involving 15 eGenes across 37 tissues (**Supplementary Table 4**).

Restricting colocalization results to those SNP-gene pairs arising from the tissue assignments provided by the TOA-classifier (at a threshold of 0.2) reduced the number of colocalizations to 133 at 32 signals, a 65% reduction overall, and a 36% reduction (from 209 at 49 signals) if considering only the subset of colocalizations that involved the four T2D-relevant tissues (Supplementary Table 5). This reduced set of TOA-filtered colocalizations retained many of the T2D effector transcripts previously reported in the literature, including those benefiting from additional chromatin conformation data^7, 8^. For example, the primary signal at the *CDC123-CAMK1D* locus (rs11257655; PPA=1.0) was classified as an islet signal (TOA=0.40) and has been previously reported to colocalize with an eQTL for *CAMK1D* expression in human islets^5, 18^. The regulatory element harboring this variant was recently shown, using promoter capture HiC, to physically interact with the *CAMK1D* promoter in human islet cells^8^. Similarly, the designation of islet signals at the *MTNR1B* (rs10830963; PPA=1.0; TOA=1.0) and *IGF2BP2* (rs150111048; PPA=0.94; TOA=0.96) loci was consistent with colocalized eQTLs implicating *MTNR1B* and *IGF2BP2* as effector genes at these loci influencing T2D risk through effects on human islet function^5, 7^.

At other signals, the integration of TOA scores with eQTL colocalization data allowed us to further resolve signals that featured multiple candidate eGenes in T2D-relevant tissues. For example, the lead SNP at the *CCND2* locus (rs76895963; PPA=1) has 16 eQTL colocalizations, involving three eGenes across 11 tissues. Of these, only two involved any of the four T2D-relevant tissues, implicating *CCND2* expression in subcutaneous adipose (CLPP=1.0) and skeletal muscle (CLPP=1.0). From a TOA perspective, this signal was classified as “shared” with high TOA scores for both islet (0.53) and adipose (0.47). This suggests that of the two colocalized eQTLs, the eQTL affecting *CCND2* expression in adipose tissue is likely to be more important to T2D pathophysiology. *CCND2* encodes cyclin D2, a signaling protein involved in cell cycle regulation and cell division. Consistent with our inference, *CCND2* was previously shown to be differentially expressed between insulin-sensitive and insulin-resistant individuals in subcutaneous adipose tissue but not in skeletal muscle^41^.

At the *CLUAP1* locus, referred to above, the lead signal (rs3751837) was classified as “shared” with comparable TOA scores across each of the four T2D-relevant tissues (0.22-0.29). Restricting to these four tissues, reduced the overall number of colocalizations (across genes and tissues) from 64 to 16. Of the remaining colocalized eQTLs, the highest colocalization posterior probability (CLPP=0.41) corresponded to an eQTL where the T2D-risk allele associates with increased expression of *TRAP1* in subcutaneous adipose (**Supplementary Table 5**). This variant is also associated with *TRAP1* expression in skeletal muscle. *TRAP1* encodes TNF Receptor Associated Protein 1, a chaperone protein that expresses ATPase activity and functions as a negative regulator of mitochondrial respiration, modulating the metabolic balance between oxidative phosphorylation and aerobic glycolysis^42^. Although *TRAP1* has not been directly implicated in T2D risk, a proteomic analysis has previously found *TRAP1* protein levels to be differentially abundant in cultured myotubes from T2D patients versus normal glucose tolerant donors^43^. Further experimental validation will be required to resolve the effector transcript(s) at this and other T2D-associated loci. However, these results, collectively, demonstrate that TOA scores can be systematically incorporated into integrative analyses to prioritise effector transcripts, particularly when there are multiple candidate genes in multiple relevant tissues.

## Discussion

We have developed a principled and extensible approach for integrative multi-omic analysis to advance the resolution of genetic mechanisms at disease-associated loci by elucidating relevant *tissues-of-action*. Existing approaches in this space have focused on characterizing the contributions of tissue- and cell-type specific regulatory features to the overall genetic architecture of the complex trait of interest (e.g. through genome-wide enrichment or heritability partitioning). However, to ensure that functional follow-up is directed to appropriate cellular systems, it is also critical to understand tissue- and cell type-specific effects at each individual signal. In line with previous work, our analyses support a prominent role for pancreatic islets in the pathogenesis of T2D, but these results also emphasize the extent to which risk-associated variants may involve shared effects across multiple tissues. Some of this tissue “sharing” was the result of incomplete resolution of causal variants at less well fine-mapped signals. However, we also found multiple examples of fine-mapped signals that overlapped regulatory elements active in multiple tissues (pointing to pleiotropic effects across tissues) as well as of loci where independent signals manifested diverse tissue-of-action profiles

A salient exemplar of these scenarios for tissue “sharing” is the *TCF7L2* locus that plays a distinguished, but as yet mechanistically-unresolved role in T2D pathogenesis and is complicated by pronounced allelic heterogeneity. The tissue-of-action for the lead signal at rs7903146 has been the subject of recent debate: early studies emphasized consequences focused on islet dysfunction whereas recent data have supported a role in adipose tissue^44, 45^. Evidence from murine studies has supported an important role for *Tcf7l2* in pancreatic *β* -cell proliferation, insulin secretion, and glucose homeostasis^46–49^. In human studies, variation at rs7903146 has been associated with chromatin accessibility and *TCF7L2* gene expression in islets^18, 44^. However, *TCF7L2* activation also regulates Wnt signaling during adipogenesis and *in vivo* deactivation of *TCF7L2* protein in mature adipocytes results in hepatic insulin resistance and systemic glucose intolerance^45^. *TCF7L2* expression was also found to be downregulated in human subjects with impaired glucose tolerance and adipocyte insulin resistance^45^. Our TOA analysis of this signal yielded a profile that is consistent with shared effects in both pancreatic islets and adipocytes that jointly contribute to T2D pathogenesis. In addition, two independent signals at this locus (rs7918400 and rs140242150) had profiles that suggest a primary mechanism of action in liver, a possibility supported by *in vivo* studies linking liver-specific perturbations of *Tcf7l2* expression in adult mice to altered hepatic glucose production and glucose production^50, 51^. Overall these data lend credence to the idea that the impact of genetic variation at this locus on T2D risk is mediated through several parallel mechanisms operating via multiple tissues. This may explain why it has such a comparatively large effect on T2D-risk in humans.

In this study, we have incorporated gene-level expression data and publicly available chromatin states based on histone ChIP-seq to determine tissues-of-action at loci associated with T2D. This scheme yielded tissue designations that were supported by validation analyses (e.g. functional fine-mapping and physiological clustering) and are consistent with previously elucidated effector mechanisms at specific loci. However, such tissue designations, though informative, constitute a first step and will undoubtedly become more refined with the increasing availability and incorporation of higher resolution datasets. In particular, our approach will benefit from more extensive genetic fine-mapping that will accompany large-scale discovery efforts involving greater samples, denser imputation reference panels, and the inclusion of more diverse populations representing underrepresented genetic ancestries.

The performance of our approach will also improve with regulome maps delineated from chromatin segmentation or hierarchical clustering analyses based on an expanded set of input features (e.g. PTM and transcription factor ChIP-seq, DNA methylation, chromatin accessibility). This allows more of the genome to be assigned to a regulatory state. For example, incorporating ATAC-seq and whole-genome bisulfite sequencing, in addition to histone PTM ChIP-seq data, into a chromatin segmentation analysis of human islets reduced the proportion of quiescent regions from 6.6% to 3.1%^19, 24^. Interestingly, islet enhancer annotations characterised by the presence of mediator binding were recently shown to exhibit a notably strong enrichment of islet-specific chromatin interactions^8^; the inclusion of such input features would help to delineate regulatory annotations that can further differentiate tissue effects. Similarly, elucidating key tissues at coding variants will benefit from long-read RNA sequencing methods that will make it possible to leverage patterns of isoform expression. Furthermore, discerning molecular features under a spectrum of biological contexts (e.g. hyperglycemia, developmental stages) will provide valuable insight into the specific conditions, within tissues-of-action, that are most relevant to individual genetic signals.

Lastly, incorporating regulatory information ascertained from single-cell approaches (e.g. scRNA-seq and snATAC-seq) will advance the resolution of *cells-of-action* against different physiological backdrops. Indeed, it may be the case that some of the tissue sharing observed in this study is reflecting cell type composition *within* tissues rather than sharing *across* tissues. The inclusion of single-cell regulome maps will help resolve this question.

The strategy presented here for integrating multi-omic information can provide valuable insight for prioritising variants and determining appropriate model systems to employ in experimental validation studies. This scheme may also enhance the construction of process-specific genetic risk scores that can identify and profile individuals with genetic burden that impacts pathophysiological processes impacting specific tissues and organ systems. Lastly, this approach can be deployed more widely across other complex diseases, especially as more tissue and cell-specific data becomes available. To support this wider use, we have implemented our method and made it openly available in an R package: Tissue of ACTion scores for Investigating Complex trait-Associated Loci (TACTICAL).

## Description of Supplemental Data

Supplemental Data include nine figures and five tables.

## Data and Code Availability

The method described in this study has been implemented in an R package titled TACTICAL (Tissue of ACTion scores for Investigating Complex trait-Associated Loci). The package can be installed from GitHub through the URL: https://github.com/Jmtorres138/TACTICAL.

## Acknowledgements

ALG is a Wellcome Trust Senior Fellow in Basic Biomedical Science. MIM was a Wellcome Senior Investigator and an NIHR Senior Investigator. This work was funded in Oxford by the Wellcome Trust (095101 [ALG], 200837 [ALG], 098381 [MIM], 106130 [ALG, MIM], 203141 (ALG, MIM], 203141 [MIM]), Medical Research Council (MR/L020149/1) [MIM, ALG], European Union Horizon 2020 Programme (T2D Systems) [ALG], and NIH (U01-DK105535; U01-DK085545) [MIM, ALG] and NIHR (NF-SI-0617-10090) [MIM]. The research was funded by the National Institute for Health Research (NIHR) Oxford Biomedical Research Centre (BRC) [ALG, MIM]. AP was supported by the Rhodes Trust, the Natural Sciences and Engineering Research Council of Canada, and the Canadian Centennial Scholarship Fund. This work was also supported by Oxford Biomedical Research Computing (BMRC) facility, a joint development between the Wellcome Centre for Human Genetics and the Big Data Institute supported by Health Data Research UK and the NIHR Oxford Biomedical Research Centre. The views expressed are those of the author and not necessarily those of the NHS, the NIHR or the Department of Health.

## Web Resources

Online resources used in this study include:

1000 Genomes Project data, http://ftp.1000genomes.ebi.ac.uk

Chromatin state maps (Varshney), https://theparkerlab.med.umich.edu

DIAGRAM website, https://www.diagram-consortium.org

Ensembl gene annotations, https://www.ensembl.org

fgwas software, https://github.com/joepickrell/fgwas

GTEx Portal website, https://gtexportal.org

LD Link, https://ldlink.nci.nih.gov

NHGRI-EBI GWAS catalogue, https://www.ebi.ac.uk/gwas

SNPsnap website, https://data.broadinstitute.org/mpg/snpsnap/

