## Supplemental_Figures for "A multi-omic integrative scheme characterizes tissues of action at loci associated with type 2 diabetes"

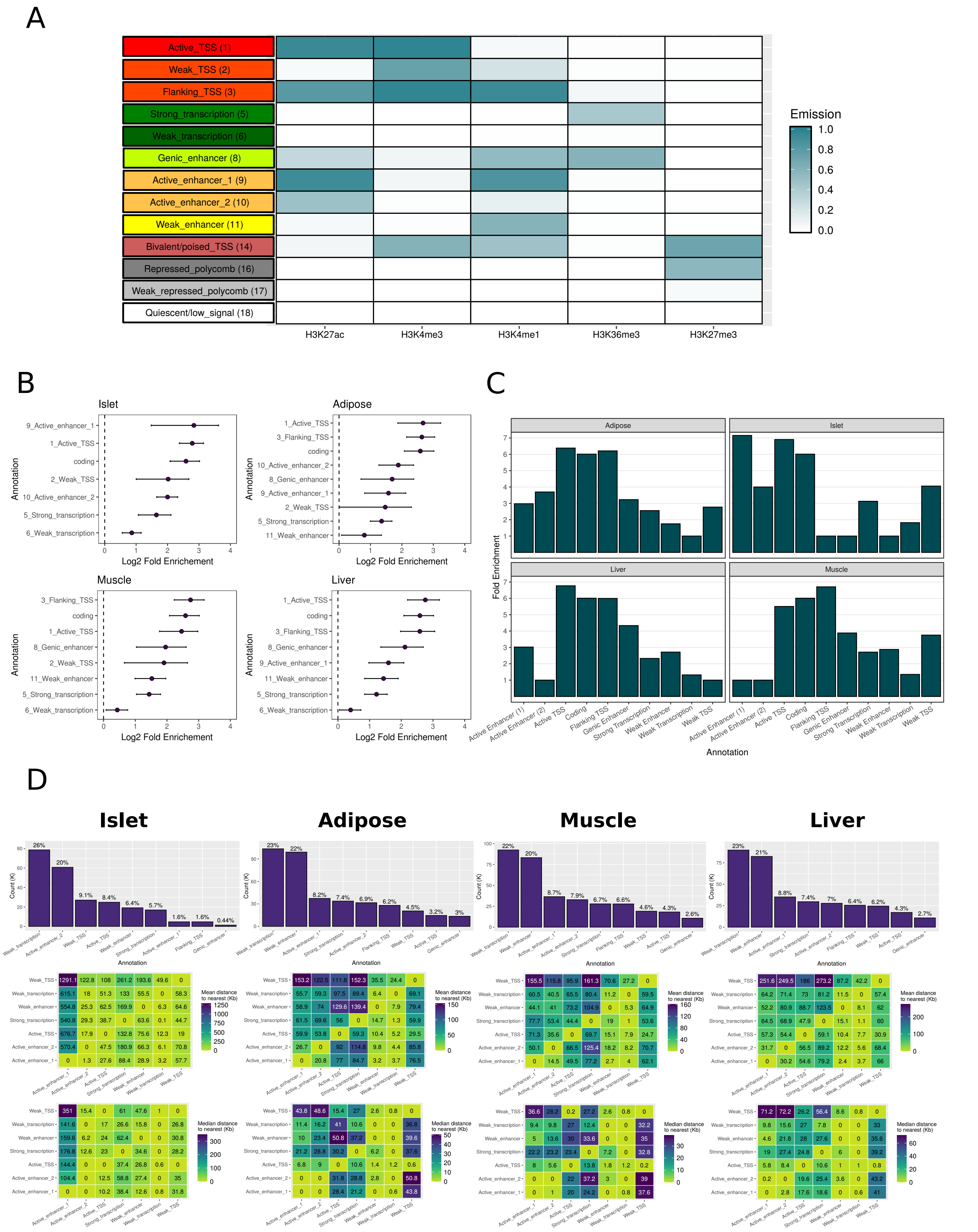
A
Active_TSS (1)
Weak_TSS (2)
Flanking_TSS (3)
Strong_transcription (5)
Weak_transcription (6)
Genic_enhancer (8)
Active_enhancer_1 (9)
Active_enhancer_2 (10)
Weak_enhancer (11)
Bivalent/poised_TSS (14)
Repressed_polycomb (16)
Weak_repressed_polycomb (17)
Quiescent/low_signal (18)
H3K27ac
H3K4me3
H3K4me1
H3K36me3
H3K27me3
Emission
1.0
0.8
0.6
0.4
0.2
0.0
B
Islet
Adipose
Muscle
Liver
Annotation
Log2 Fold Enrichement
C
Adipose
Islet
Liver
Muscle
Fold Enrichment
Annotation
Active Enhancer (1)
Active Enhancer (2)
Active TSS
Coding
Flanking TSS
Genic Enhancer
Strong Transcription
Weak Enhancer
Weak Transcription
Weak TSS
D
Islet
Adipose
Muscle
Liver
Count (K)
Annotation
Mean distance to nearest (Kb)
Median distance to nearest (Kb)

**Supplementary Figure 1. Epigenomic annotations are enriched for variants associated with type 2 diabetes. A)** ChromHMM emission values from Varshney et al. 2017. are indicated for each chromatin state annotation. **B)** Chromatin state annotations were assessed for genome-wide enrichment of T2D-associated SNPs using the program *fgwas*. $\text{Log}_2$ fold enrichment values and 95% confidence bounds are shown for each chromatin state annotation meeting statistical significance **C)** The mean fold enrichment values for each tissue annotation were used as weights for scaling the relative contribution of each annotation vector in calculating TOA scores. **D)** The count distribution of epigenomic annotations is shown for each tissue along with heat plots showing the pairwise mean and median distances to the nearest annotation in kilobases (Kb).

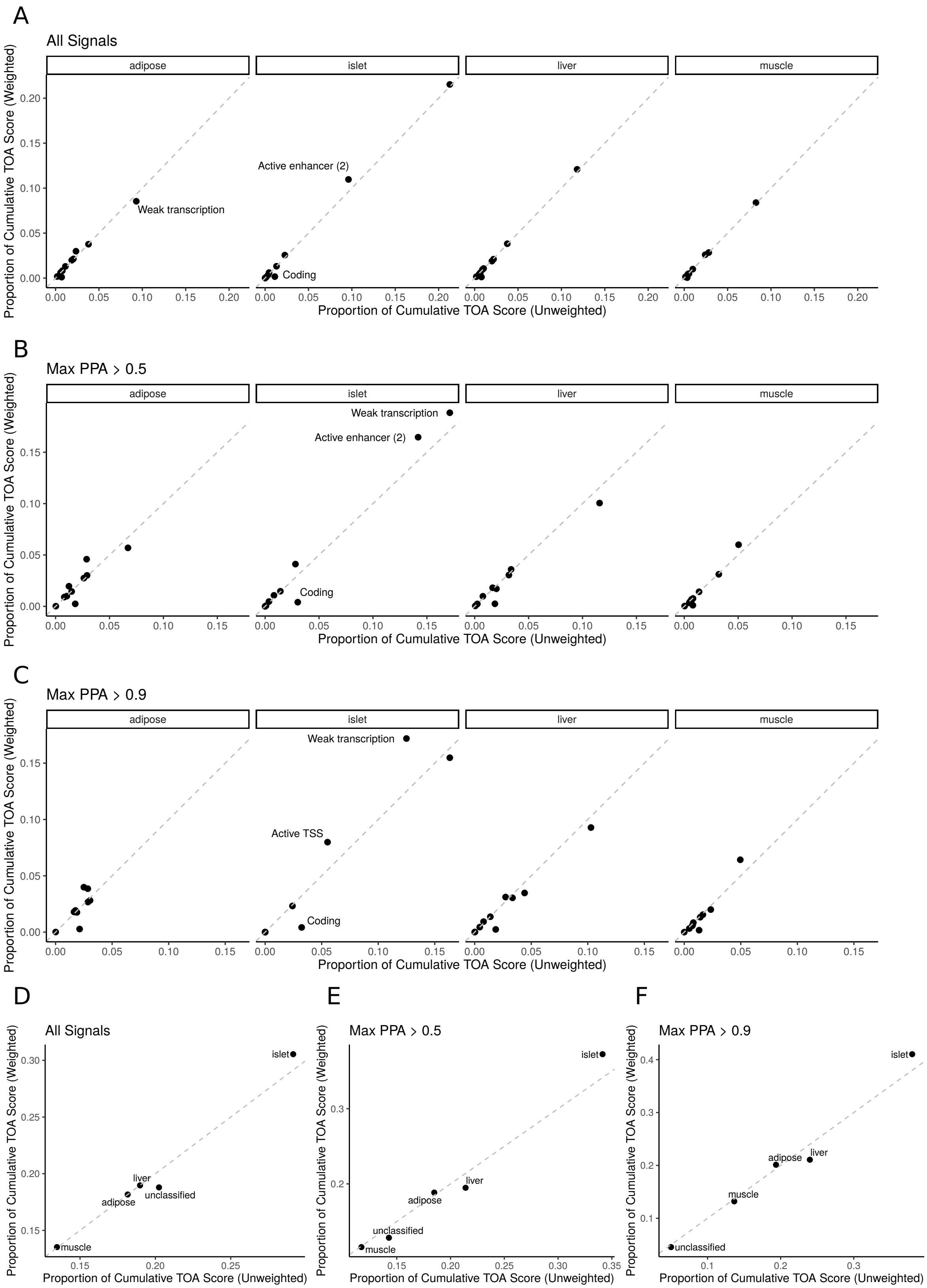
A
All Signals
adipose
islet
liver
muscle
Weak transcription
Active enhancer (2)
Coding
Proportion of Cumulative TOA Score (Weighted)
Proportion of Cumulative TOA Score (Unweighted)
B
Max PPA > 0.5
Weak transcription
Active enhancer (2)
Coding
C
Max PPA > 0.9
Weak transcription
Active TSS
Coding
D
All Signals
islet
liver
adipose
unclassified
muscle
E
Max PPA > 0.5
islet
liver
adipose
unclassified
muscle
F
Max PPA > 0.9
islet
adipose
liver
muscle
unclassified

**Supplementary Figure 2. Comparison of GWAS effect size weighted and unweighted TOA scores.** TOA scores weighted by $\frac{|\beta|}{SE}$, corresponding to the absolute value of the GWAS effect size divided by standard error of the sentinel SNP corresponding to each signal, are compared with the unweighted TOA scores. The proportion of cumulative score for unweighted (x-axis) and weighted (y-axis) TOA scores are shown for each annotation group in each tissue and is shown for **(A)** all signals, **(B)** all signals with maximum credible set PPA $> 0.5$, and **(C)** all signals with maximum credible set PPA $> 0.9$. The three annotation groups with the greatest differences between weighted and unweighted TOA scores, across all tissues, are indicated. The proportions of cumulative TOA scores combined across annotation groups for each tissue are shown for **(D)** all signals, **(E)** all signals with maximum credible set PPA $> 0.5$, and **(F)** all signals with maximum credible set PPA $> 0.9$.

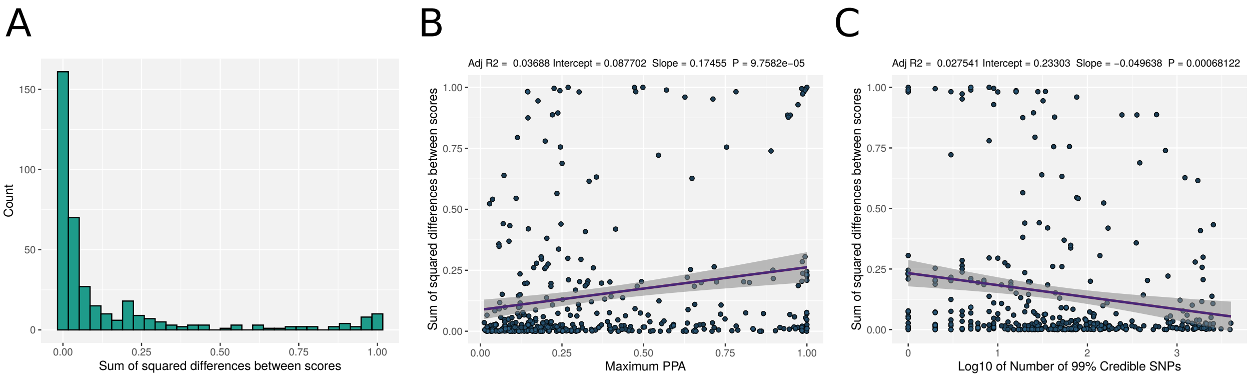


**Supplementary Figure 3. The relationship between credible set resolution and tissue specificity.** **A)** Histogram of sum of squared differences (SSD) between tissue-of-action scores for islet, subcutaneous adipose, skeletal muscle, and liver across all 380 fine-mapped signals. **B)** Scatter plot of maximum 99% credible set PPA (x-axis) and SSD values (y-axis) is shown for all signals with regression line. **C)** Scatter plot of base 10 logarithm of number of 99% credible SNPs (x-axis) and SSD values (y-axis) is shown for all signals with regression line.

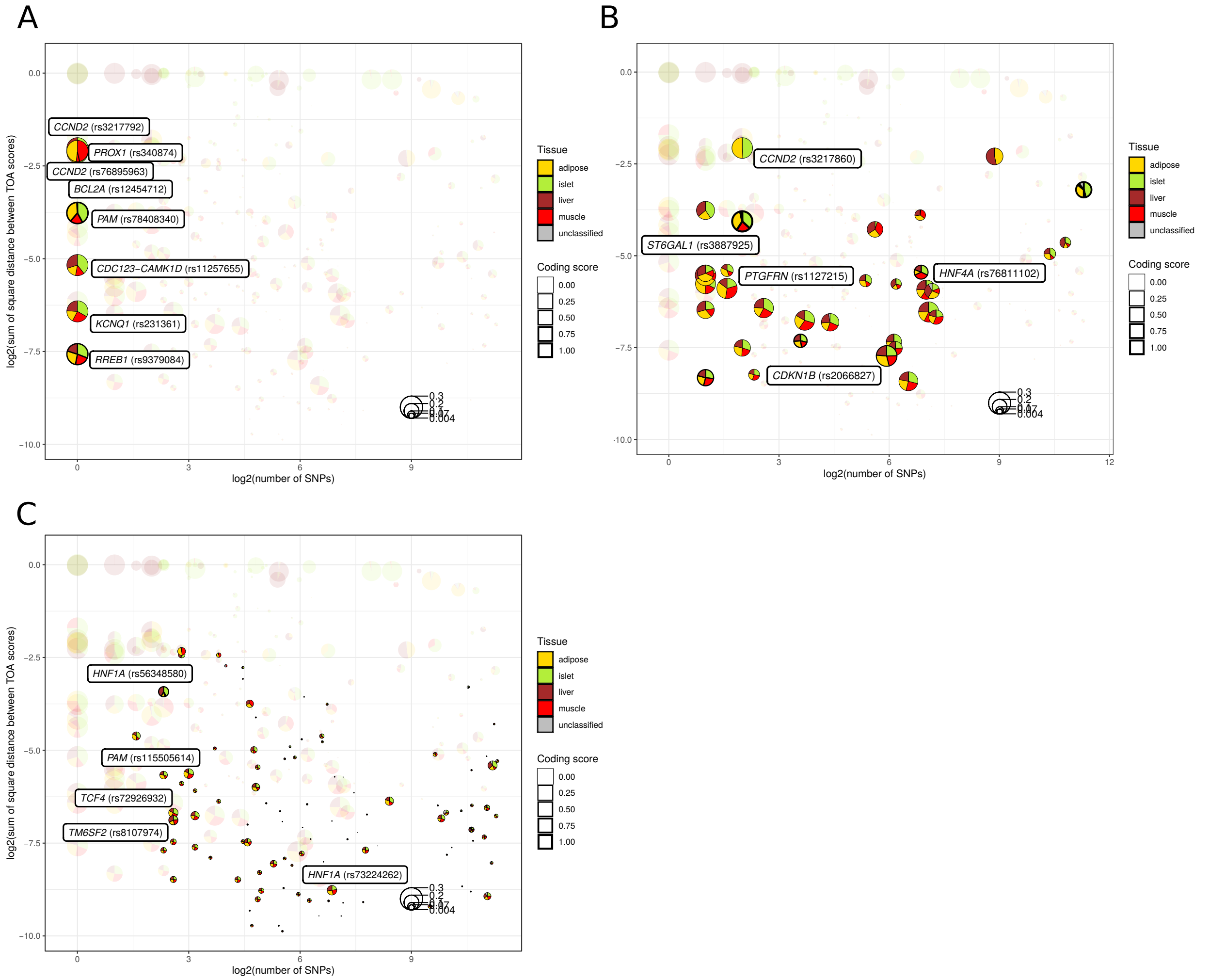


**Supplementary Figure 4. Fine-mapping resolution and TOA score diversity at shared signals.** **A)** Signals that have been fine-mapped to a single credible variant. **B)** Signals with more than one credible variant but maximum PPA > 0.5. **C)** Signals with more than one credible variant and maximum PPA ≤ 0.5. $\text{Log}_2$ of the number of credible SNPs for each fine-mapped signal is shown on the x-axis and the $\log_2$ value of the sum of square differences between TOA scores for each signal is shown on the y-axis (i.e. higher values on the y-axis correspond to greater tissue "specificity"). The profile of TOA scores are indicated within pie charts where the diameter of each circle corresponds to the maximum PPA for the credible set. The line thickness for each circle indicates a coding score for each credible set (i.e. the proportion of cumulative PPA attributable to coding variants).

A

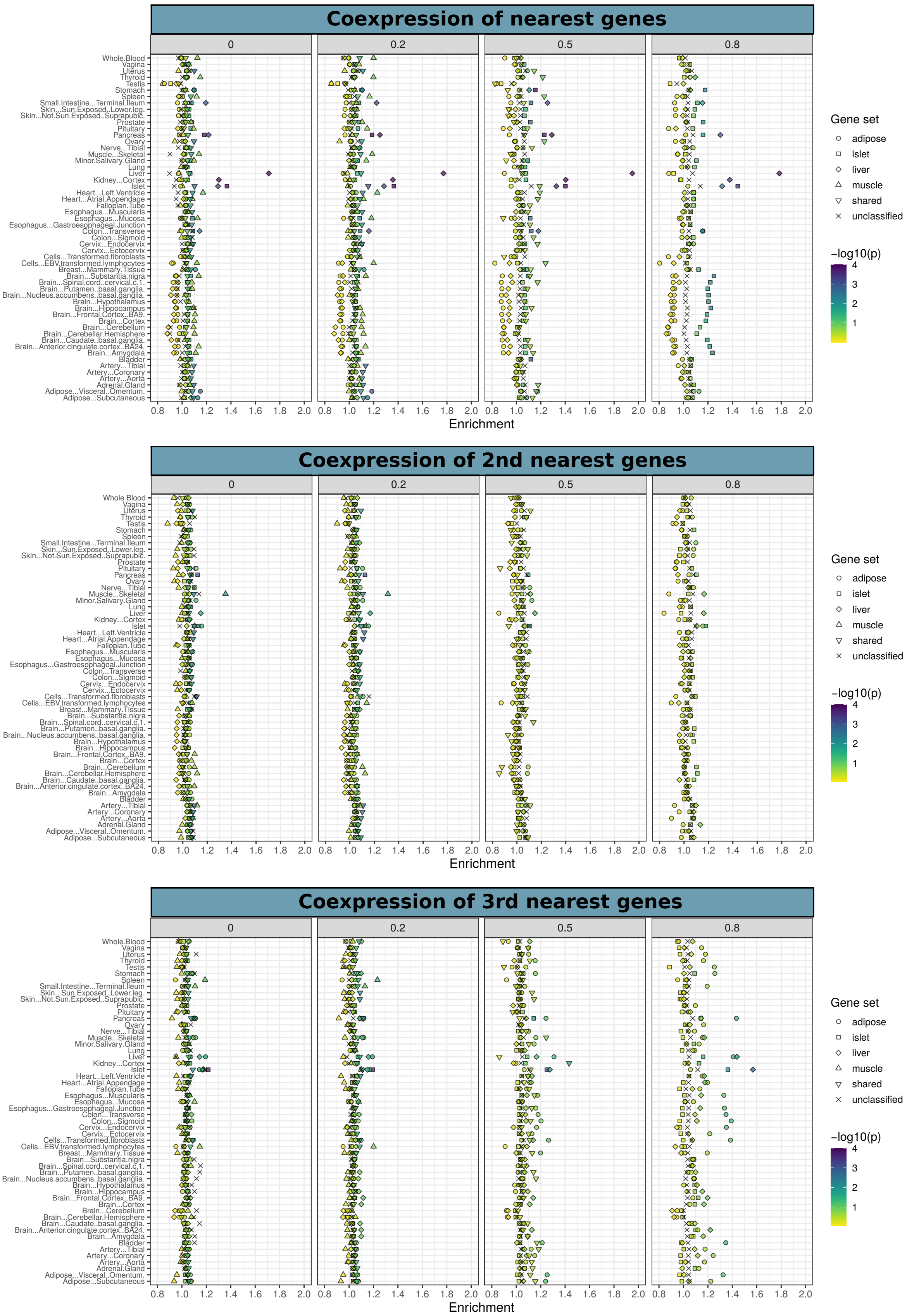

Coexpression of nearest genes
Coexpression of 2nd nearest genes
Coexpression of 3rd nearest genes
0
0.2
0.5
0.8
Tissue
Whole.Blood
Vagina
Uterus
Thyroid
Testis
Stomach
Spleen
Small.Intestine...Terminal.Ileum
Skin...Sun.Exposed..Lower.leg.
Skin...Not.Sun.Exposed..Suprapubic.
Prostate
Pituitary
Pancreas
Ovary
Nerve...Tibial
Muscle...Skeletal
Minor.Salivary.Gland
Lung
Liver
Kidney...Cortex
Islet
Heart...Left.Ventricle
Heart...Atrial.Appendage
Fallopian.Tube
Esophagus...Muscularis
Esophagus...Mucosa
Esophagus...Gastroesophageal.Junction
Colon...Transverse
Colon...Sigmoid
Cervix...Endocervix
Cervix...Ectocervix
Cells...Transformed.fibroblasts
Cells...EBV.transformed.lymphocytes
Breast...Mammary.Tissue
Brain...Substantia.nigra
Brain...Spinal.cord..cervical.c.1.
Brain...Putamen..basal.ganglia.
Brain...Nucleus.accumbens..basal.ganglia.
Brain...Hypothalamus
Brain...Hippocampus
Brain...Frontal.Cortex..BA9.
Brain...Cortex
Brain...Cerebellum
Brain...Cerebellar.Hemisphere
Brain...Caudate..basal.ganglia.
Brain...Anterior.cingulate.cortex..BA24.
Brain...Amygdala
Bladder
Artery...Tibial
Artery...Coronary
Artery...Aorta
Adrenal.Gland
Adipose...Visceral..Omentum.
Adipose...Subcutaneous
0.8 1.0 1.2 1.4 1.6 1.8 2.0
Enrichment
Gene set
adipose
islet
liver
muscle
shared
unclassified
−log10(p)
4
3
2
1


B

C

**Supplementary Figure 5. Coexpression of proximal genes to tissue-assigned signals. A)** Significance of coexpression of nearest genes annotated to sets of tissue-assigned signals across TOA score stringency thresholds. Coexpression for sets of second nearest and third nearest genes are shown in **B)** and **C)**, respectively. Results are shown for human islet and 53 tissues from the Genotype-Tissue Expression Project (GTEx). Shape indicates the tissue to which the set of signals were assigned.

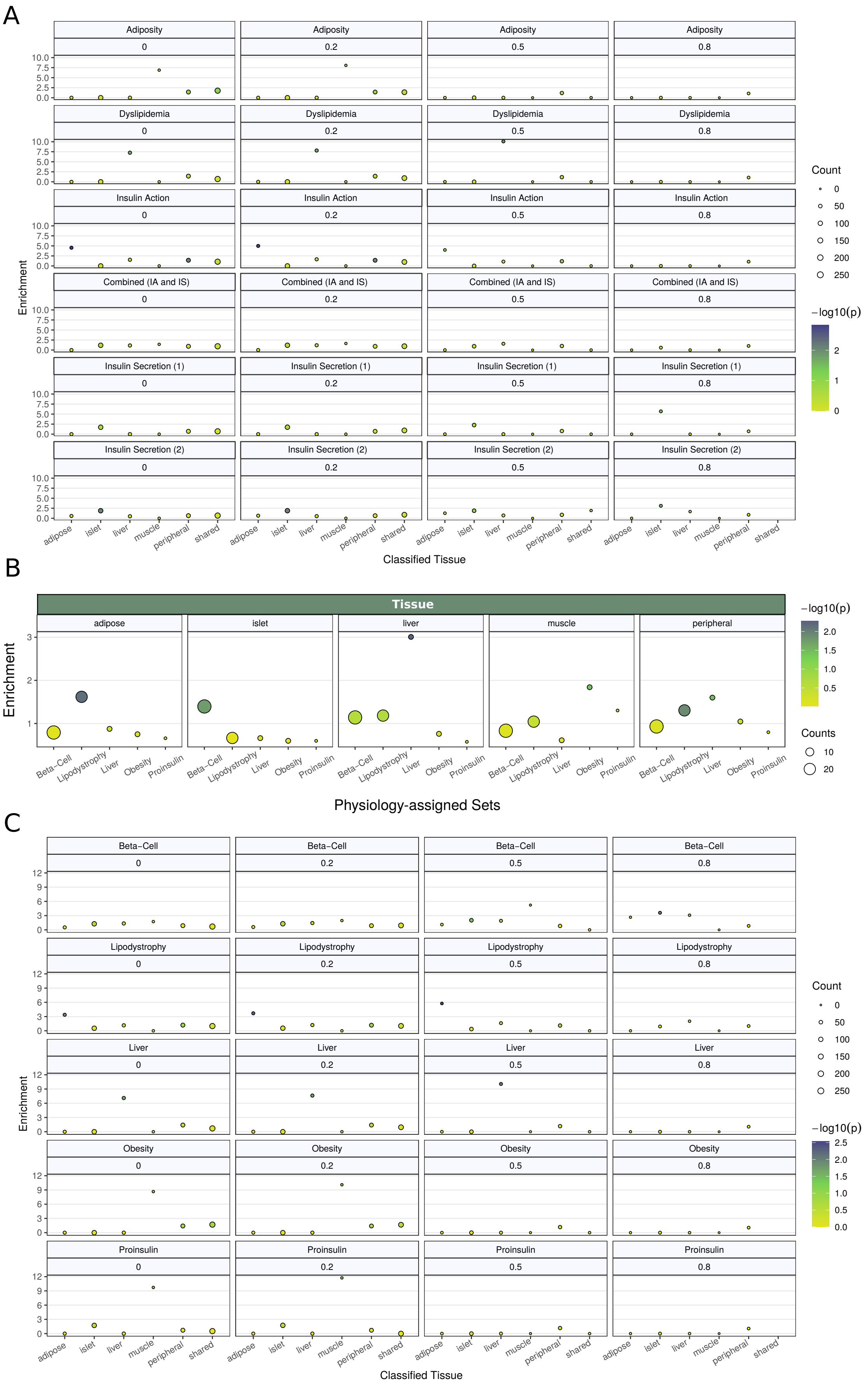
A
Adiposity
Dyslipidemia
Insulin Action
Combined (IA and IS)
Insulin Secretion (1)
Insulin Secretion (2)
0
0.2
0.5
0.8
Enrichment
adipose
islet
liver
muscle
peripheral
shared
Classified Tissue
Count
−log10(p)
B
Tissue
adipose
islet
liver
muscle
peripheral
Beta−Cell
Lipodystrophy
Liver
Obesity
Proinsulin
Physiology-assigned Sets
Counts
C
Beta−Cell
Lipodystrophy
Liver
Obesity
Proinsulin

**Supplementary Figure 6. Comparison of sets of epigenomic- and physiology-assigned signals.** **A)** Enrichment of 82 physiology-assigned signals, from six physiology groups, among sets of epigenomic-assigned signals based on TOA scores. Size corresponds to the number of physiology signals among each set of tissue-assigned signals. **B)** Comparison of TOA scores - as enrichments of mean observed value over a sampled mean - among sets of physiology-assigned signals from Udler et al. 2018. The size corresponds to the number assigned signals in each physiology group. **C)** Enrichment of 62 physiology-assigned signals from Udler et al. 2018., from five physiology groups, among sets of epigenomic-assigned signals based on TOA scores. Size corresponds to the number of physiology signals among each set of tissue-assigned signals.

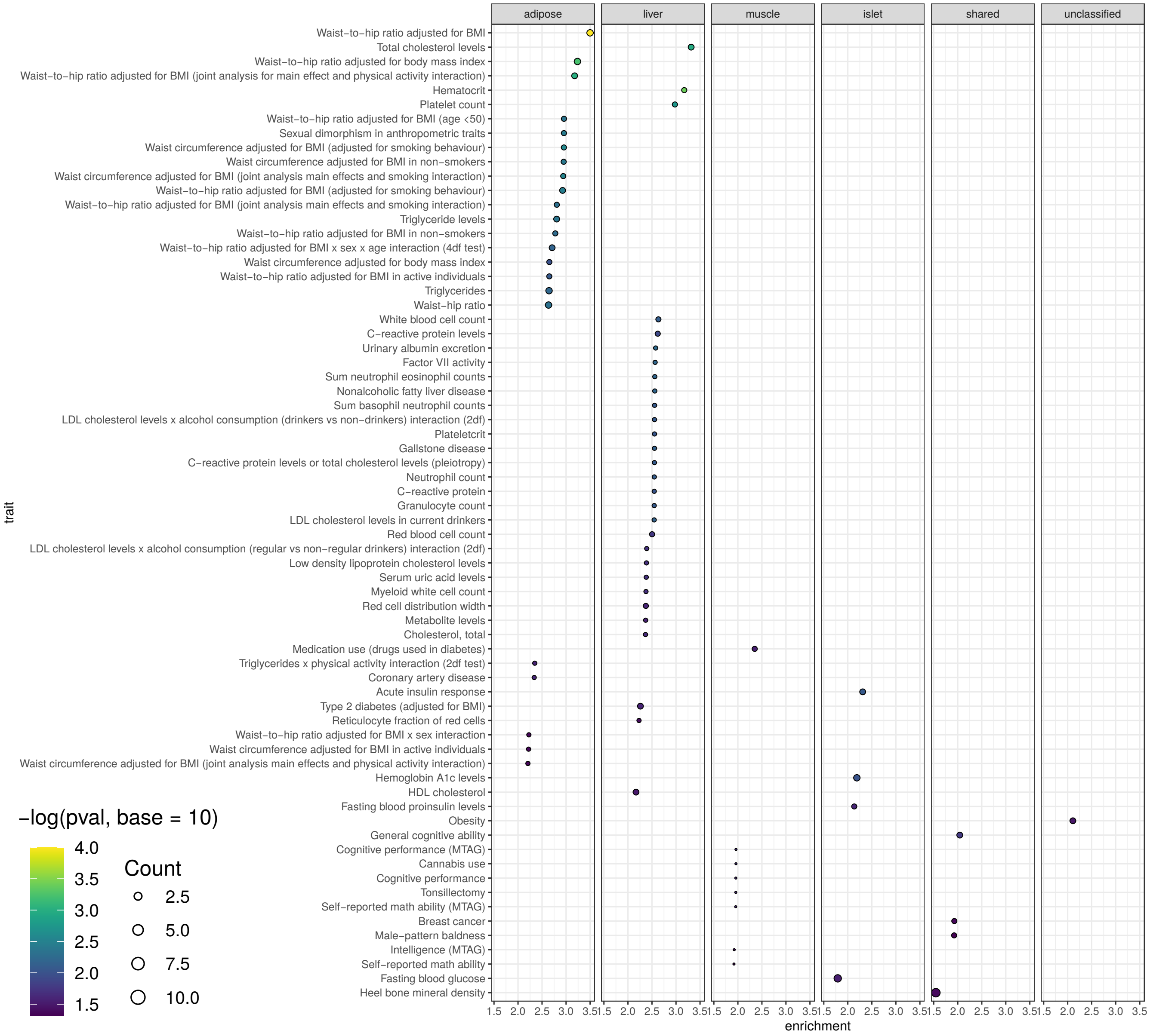


**Supplementary Figure 7. Enrichment of trait-associated SNPs.** The results from an enrichment test are shown each trait from the NHGRI-EBI GWAS catalogue where trait associated SNPs were enriched among sets of tissue-assigned signals at a p-value $\leq 0.05$. All results correspond to signals classified at the 0.2 TOA score threshold. Color corresponds to the negative $\log_{10}$ p-value and the text size corresponds to the number of SNPs that overlap lead SNPs (through identity or LD proxy at r2 $\geq 0.8$) for each tissue-assigned signal.

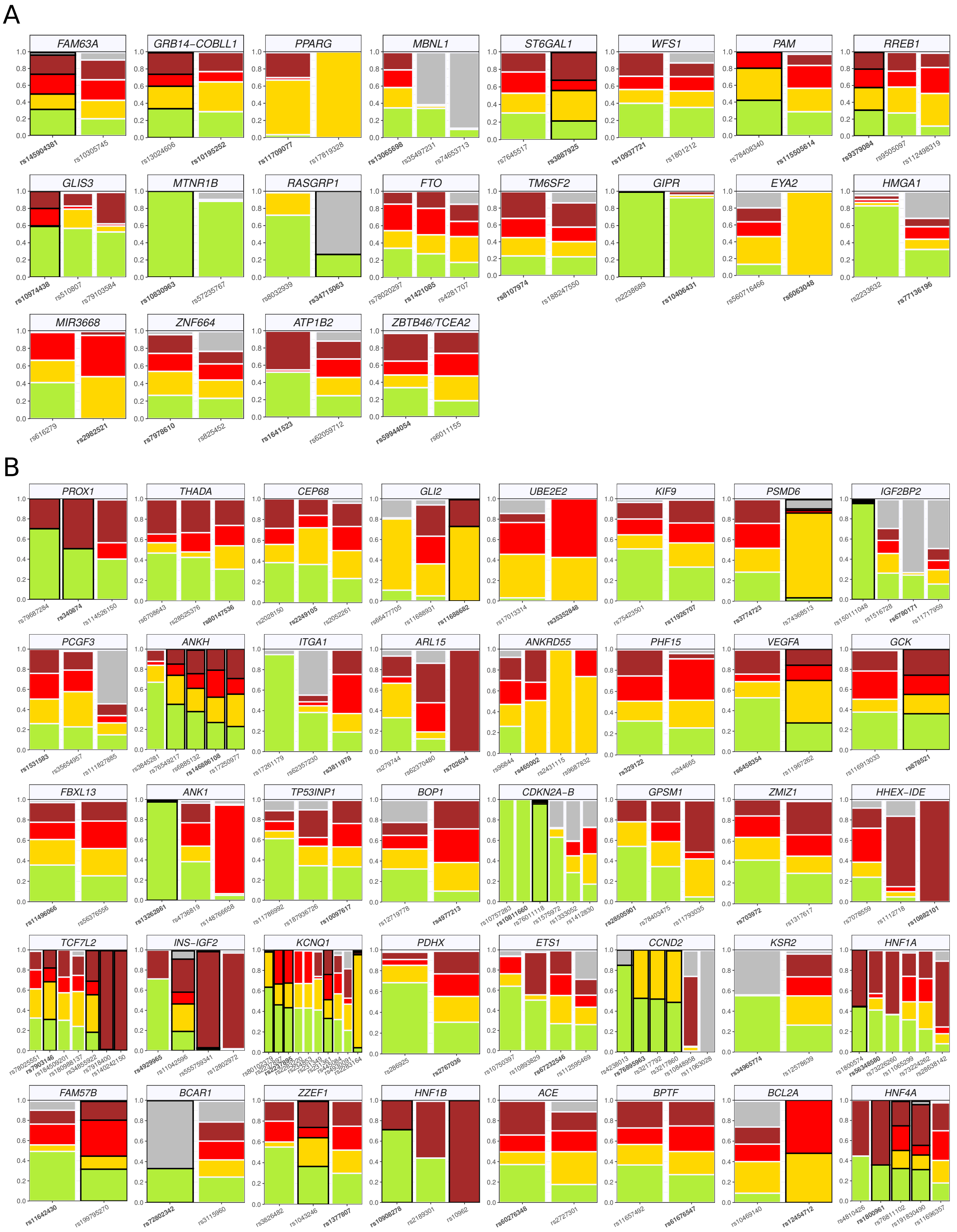


**Supplementary Figure 8.** **Multiple tissues implicated by epigenomic scores at heterogenous loci.** **A)** Profile of TOA scores for the 20 loci with all signals receiving identical tissue assignments at the 0.2 stringency threshold **B)** Profile of TOA scores for the 40 loci with all signals receiving distinct tissue assignments at the 0.2 threshold.

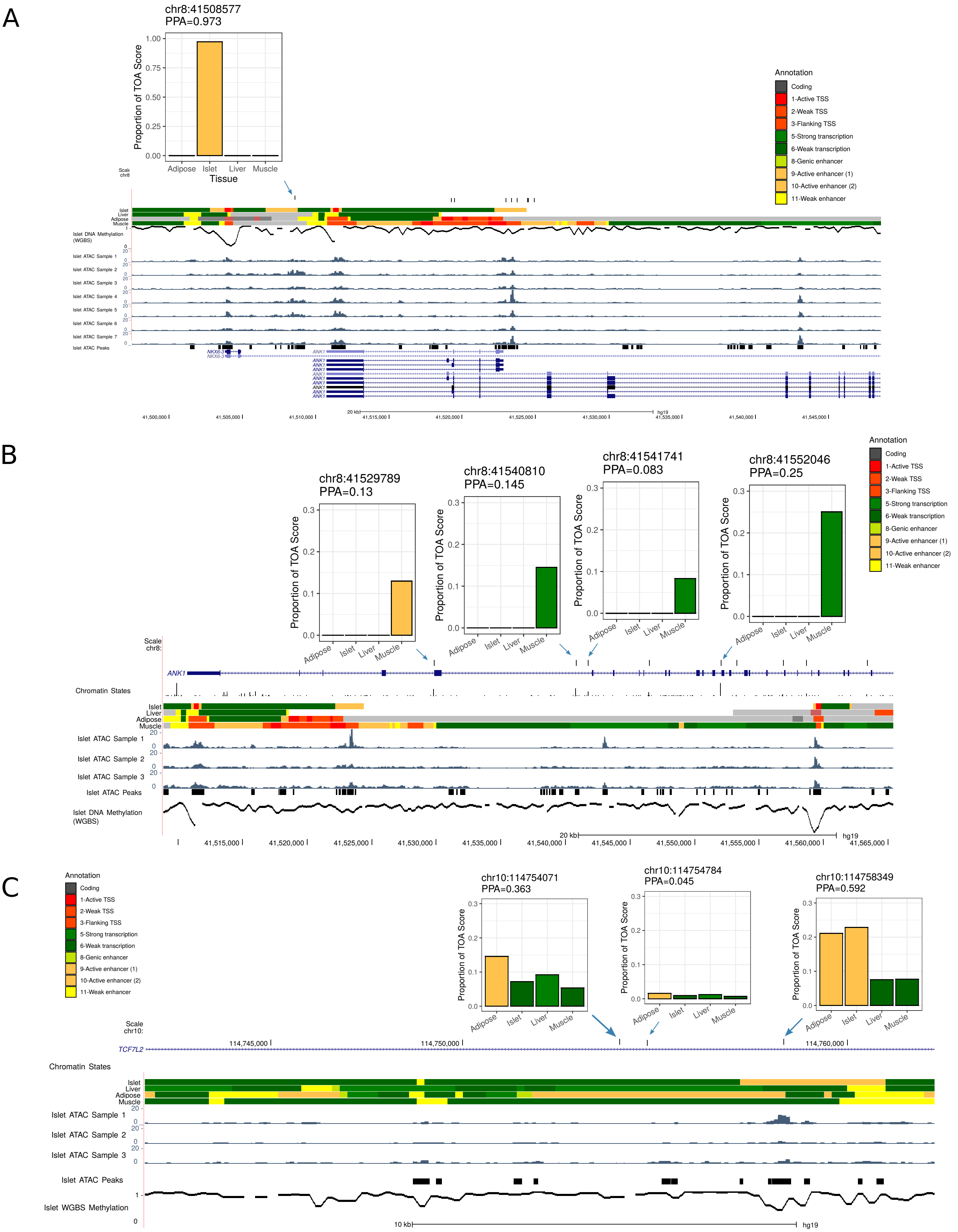
A
chr8:41508577
PPA=0.973
Proportion of TOA Score
Adipose
Islet
Liver
Muscle
Tissue
Annotation
Coding
1-Active TSS
2-Weak TSS
3-Flanking TSS
5-Strong transcription
6-Weak transcription
8-Genic enhancer
9-Active enhancer (1)
10-Active enhancer (2)
11-Weak enhancer
Islet DNA Methylation (WGBS)
Islet ATAC Sample 1
Islet ATAC Sample 2
Islet ATAC Sample 3
Islet ATAC Sample 4
Islet ATAC Sample 5
Islet ATAC Sample 6
Islet ATAC Sample 7
Islet ATAC Peaks
NKX6-3
ANK1
B
chr8:41529789
PPA=0.13
chr8:41540810
PPA=0.145
chr8:41541741
PPA=0.083
chr8:41552046
PPA=0.25
ANK1
Chromatin States
Islet ATAC Sample 1
Islet ATAC Sample 2
Islet ATAC Sample 3
Islet ATAC Peaks
Islet DNA Methylation (WGBS)
C
chr10:114754071
PPA=0.363
chr10:114754784
PPA=0.045
chr10:114758349
PPA=0.592
TCF7L2
Chromatin States
Islet ATAC Sample 1
Islet ATAC Sample 2
Islet ATAC Sample 3
Islet ATAC Peaks
Islet WGBS Methylation
10 kb
hg19

**Supplementary Figure 9. Epigenomic profiles of fine-mapped signals at T2D-associated loci. A)** Islet-assigned and primary signal (rs13262861) at the *ANK1* locus. **B)** Muscle-assigned signal at the *ANK1* locus (rs148766658). **C)** Islet-assigned and primary signal (rs7903146) at the *TCF7L2* locus. For each credible SNP, the PPA value attributable to each tissue annotation is shown along with its position on chromosome 10 (genome build hg19). Chromatin state maps for islet, adipose, muscle, and liver tissue from Varshney et al. 2017. are shown along with ATAC-seq tracks for representative islet samples, called ATAC-seq peaks from a set of islet ATAC samples (n=17), and DNA methylation (whole genome bisulfite sequencing) in human islets from Thurner et al. 2018.
